# Temporal evolution of single-cell transcriptomes of *Drosophila* olfactory projection neurons

**DOI:** 10.1101/2020.09.24.312397

**Authors:** Qijing Xie, Maria Brbic, Felix Horns, Sai Saroja Kolluru, Bob Jones, Jiefu Li, Anay Reddy, Anthony Xie, Sayeh Kohani, Zhuoran Li, Colleen McLaughlin, Tongchao Li, Chuanyun Xu, David Vacek, David J. Luginbuhl, Jure Leskovec, Stephen R. Quake, Liqun Luo, Hongjie Li

## Abstract

Neurons undergo substantial morphological and functional changes during development to form precise synaptic connections and acquire specific physiological features. What are the underlying transcriptomic bases? Here, we obtained the single-cell transcriptomes of *Drosophila* olfactory projection neurons (PNs) at four developmental stages. We decoded the identity of 21 transcriptomic clusters corresponding to 20 PN types and developed methods to match transcriptomic clusters representing the same PN type across development. We discovered that PN transcriptomes reflect unique biological processes unfolding at each stage—neurite growth and pruning during metamorphosis at an early pupal stage; peaked transcriptomic diversity during olfactory circuit assembly at mid-pupal stages; and neuronal signaling in adults. At early developmental stages, PN types with adjacent birth order share similar transcriptomes. Together, our work reveals principles of cellular diversity during brain development and provides a resource for future studies of neural development in PNs and other neuronal types.

## Introduction

Cell-type diversity and connection specificity between neurons are the basis of accurate information processing underlying all nervous system functions. The precise assembly of neural circuits involves multiple highly regulated steps. First, neurons are born from their progenitors and acquire unique fates through a combination of (1) intrinsic mechanisms, such as lineage, birth order, and birth timing; (2) extrinsic mechanisms, such as lateral inhibition and extracellular induction, and (3) developmental stochasticity in some cases (Jan & Jan, 1994; Johnston & Desplan, 2010; Kohwi & Doe, 2013; Holguera & Desplan, 2018; Li et al., 2018). During wiring, neurons extend their neurites to a coarse targeting region, elaborate their terminal structures, select pre- and post-synaptic partners, and finally form synaptic connections (Sanes & Yamagata, 2009; Jan & Jan, 2010; Kolodkin & Tessier-Lavigne, 2011; Sanes & Zipursky, 2020). Studies from the past few decades have uncovered many molecules and mechanisms that regulate each of these developmental processes.

The development of *Drosophila* olfactory projection neurons (PNs) has been extensively studied (Jefferis et al., 2004; Hong & Luo, 2014). PNs are the second-order olfactory neurons that receive organized input from olfactory receptor neurons (ORNs) at ∼50 stereotyped and individually identifiable glomeruli in the antennal lobe, and carry olfactory information to higher brain regions (Vosshall & Stocker, 2007; Wilson, 2013) (Figure 1A). Different types of PNs send their dendrites to a single glomerulus or multiple glomeruli (Marin et al., 2002; Lai et al., 2008; Yu et al., 2010; Tanaka et al., 2012; Bates et al., 2020). PNs are derived from three separate neuroblast lineages—anterodorsal, lateral, and ventral lineages, corresponding to their cell bodies’ positions relative to the antennal lobe (Jefferis et al., 2001). PNs produced from the anterodorsal and lateral lineages (adPNs and lPNs) are cholinergic excitatory neurons. The fate of uniglomerular excitatory PN types, defined by their glomerular targets, is predetermined by their lineage and birth order (Jefferis et al., 2001; Yu et al., 2010; Lin et al., 2012). PNs produced from the ventral lineage (vPNs), on the other hand, are GABAergic inhibitory neurons (Jefferis et al., 2007; Liang et al., 2013; Parnas et al., 2013). The connectivity and physiology of PNs have also been systematically studied (Bhandawat et al., 2007; Jeanne et al., 2018; Bates et al., 2020).

**Figure 1.**
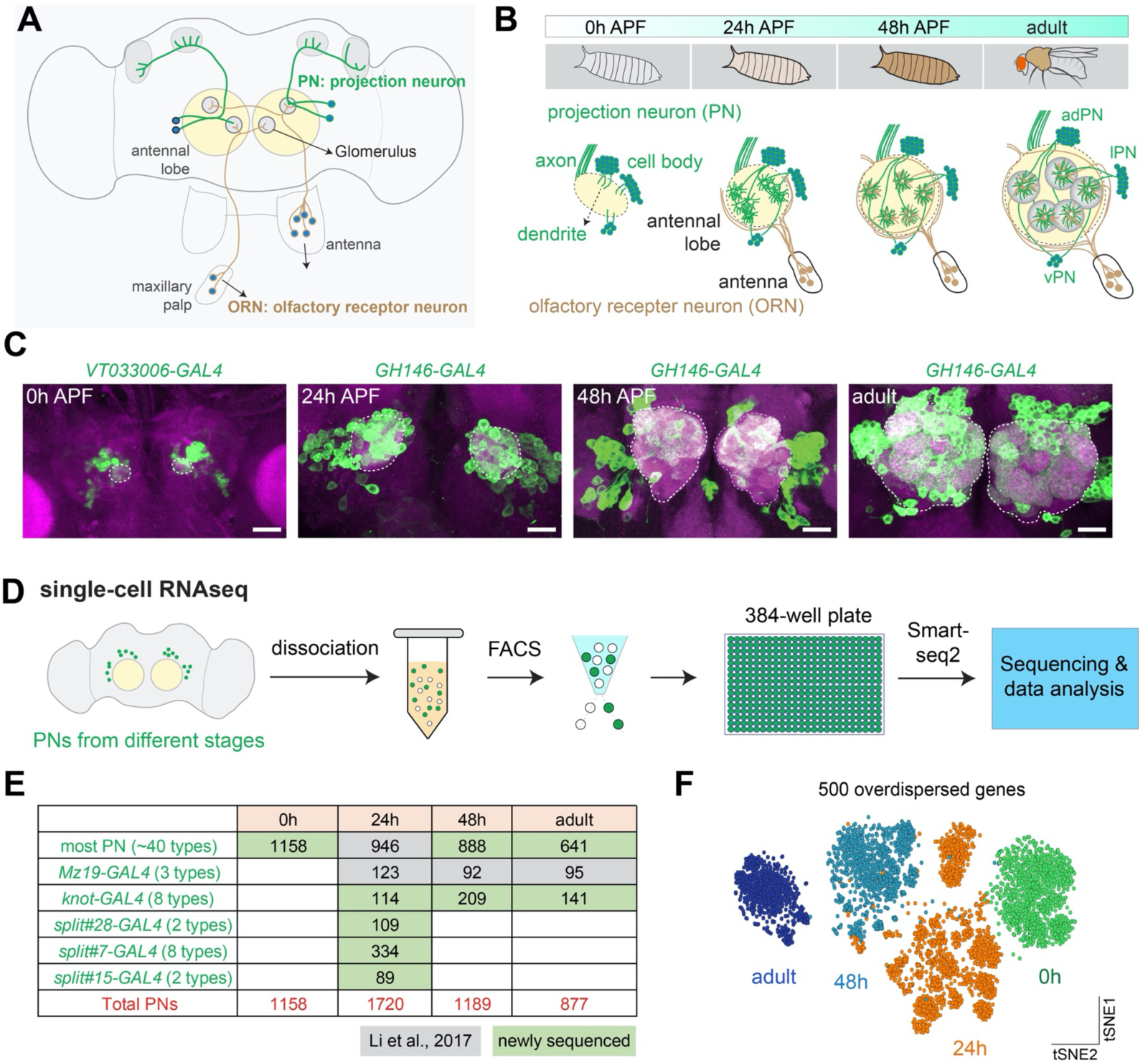
Overview of single-cell transcriptomic profiling of *Drosophila* olfactory projection neurons (PNs). **(A)** Schematic of the adult *Drosophila* olfactory system. 50 types of olfactory receptor neurons (ORNs) form one-to-one synaptic connections with 50 types of excitatory PNs at 50 glomeruli in the antennal lobe. Illustrated are two types each of ORNs (brown) and PNs (green), as well as two glomeruli to which their axons and dendrites target. **(B)** Schematic of the developmental process of the adult *Drosophila* olfactory system. The ∼50 types of excitatory PNs are from either anterodorsal (adPN) or lateral (lPN) neuroblast lineages. PNs with cell body on the ventral side are inhibitory ventral PNs (vPNs). **(C)** Representative confocal images of PNs from four different developmental stages, 0h APF, 24h APF, 48h AFP, and adult. APF: after puparium formation. Images are shown as maximum z-projections of confocal stacks. Antenna lobe is outlined. Scale bars, 40 μm. **(D)** Workflow of the single-cell RNA sequencing using plate-based SMART-seq2. FACS: fluorescence-activated cell sorting. **(E)** Summary of the number of high-quality PNs sequenced at each timepoint and driver lines used. Most PNs refer to PNs sequenced using either *GH146-GAL4* or *VT033006-GAL4*. **(F)** Visualization of all sequenced PNs from four different developmental stages using tSNE plot. Dimensionality reduction was performed using the top 500 overdispersed genes identified from all sequenced PNs.

Despite the fact that PNs are among the most well-characterized cell types in all nervous systems, their genome-wide gene expression changes across different developmental stages with cell-type specificity are still unknown. This information can help us obtain a more complete picture of both known and unexplored pathways underlying neural development and function. Recently, the advent of single-cell RNA sequencing (scRNA-seq) has paved the way towards obtaining such data (Li et al., 2017; Kalish et al., 2018; Zhong et al., 2018; Li, 2020). Here, we profiled and analyzed the single-cell transcriptomes of most uniglomerular excitatory PNs. We identified the correspondence between two-thirds of transcriptomes and PN types at one stage, and developed methods to reliably match transcriptomic clusters corresponding to the same types of PNs across different stages. We discovered that PN transcriptomes exhibit unique characteristics at different stages, including birth-order, neurite pruning, wiring specificity, and neuronal signaling.

## Results

### Single-cell transcriptomic profiling of *Drosophila* PNs at four developmental stages

The development of PNs follows the coordinated steps previously described. 18 out of 40 types of adPNs are born embryonically and participate in the larval olfactory system. Then, during the larval stage, the rest of adPNs and all lPNs are born (Jefferis et al., 2001; Marin et al., 2005; Yu et al., 2010; Lin et al., 2012). During metamorphosis following puparium formation, embryonically born PNs first prune terminal branches of dendrites and axons, and then re-extend their dendrites into the future adult antennal lobe, and axons into the mushroom body and lateral horn following the neurites of larval-born PNs (Marin et al., 2005). From 0 to 24 hours after puparium formation (APF), PNs extend their dendrites into the developing antennal lobe and occupy restricted regions. ORN axons begin to invade antennal lobe at ∼24 hours APF. PN dendrites and ORN axons then match with their respective partners beginning at ∼30 hours APF and establish discrete glomerular compartments at ∼48 hours APF. Thereafter, they expand their terminal branches, build synaptic connections, and finally form mature adult olfactory circuits (Jefferis et al., 2004) (Figure 1B).

To better understand the molecular mechanisms that control these dynamic developmental processes underlying neural circuit assembly, we performed scRNA-seq of PNs from 4 different developmental stages: 0–6 hours APF, 24–30 hours APF, 48–54 hours APF, and 1–5 days adult (hereafter 0, 24, 48h APF and adult) (Figure 1C). We used *GH146-GAL4* (Stocker et al., 1997) to drive *UAS-mCD8-GFP* (Lee et al., 1999) expression in most PNs at 24h, 48h, and adult, which labels ∼90 of the estimated 150 PNs in each hemisphere, covering ∼40 of the 50 PN types. At 0h APF, *GH146-GAL4* also labels cells in the optic lobes (Figure 1—figure supplement 1A), which are inseparable from the central brain by dissection. Therefore, we used *VT033006-GAL4* to label PNs at 0h APF (Figure 1C and Figure 1—figure supplement 1B) (Tirian & Dickson, 2017). *VT033006-GAL4* labels most PNs from the anterodrosal and lateral lineage, but not PNs from the ventral lineage or anterior paired lateral (APL) neurons like *GH146-GAL4*. It is expressed in ∼95 cells that innervate ∼44 glomeruli which largely overlap with PNs labeled by *GH146-GAL4* (Inada et al., 2017; Elkahlah et al., 2020). In addition to PNs labeled by *GH146-GAL4* and *VT033006-GAL4* (we will refer them as ‘most PNs’ hereafter), we have collected single-cell transcriptomic data using drivers that only label a small number of PN types for mapping the transcriptomic clusters to anatomically defined PN types.

For scRNA-seq, fly brains with a unique set of PN types labeled using different drivers at each developmental stage were dissected and dissociated into single-cell suspensions. GFP+ cells were sorted into 384-well plates by fluorescence-activated cell sorting (FACS), and sequenced using SMART-seq2 (Picelli et al., 2014) (Figure 1D) to a depth of ∼1 million reads per cell (Figure 1–figure supplement 1C). On average ∼3000 genes were detected per cell (Figure 1–figure supplement 1D), and after quality filtering (see Methods), we obtained ∼3700 high quality PNs in addition to the previously sequenced ∼1200 PNs (Li et al., 2017), yielding ∼5900 PN cells for analysis in this study (Figure 1E). All analyzed PNs express high levels of neuronal markers but not glial markers, confirming the specificity of sequenced cells (Figure 1–figure supplement 1E). Unbiased clustering using overdispersed genes from all PNs readily separates them into different groups according to their stage (Figure 1F), suggesting that gene expression changes across these four developmental stages represent a principal difference in their single-cell transcriptomes.

### Decoding the glomerular identity of transcriptomic clusters by sequencing subsets of PNs at 24h APF

PNs labeled by *GH146-GAL4* at 24h APF form ∼30 distinct transcriptomic clusters. We previously matched 6 of these transcriptomic clusters to specific anatomically and functionally defined PN types (Li et al., 2017), hereafter referred to as “decoding transcriptomic identity.” Unlike ORNs, whose identities can be decoded using uniquely expressed olfactory receptors (Li et al., 2020a), PNs lack known type-specific markers. Instead, PN types are mostly specified by combinatorial expression of several genes (Li et al., 2017), making it more challenging to decode their transcriptomic identities.

To circumvent these challenges and decode the transcriptomic identities of more types of PNs, we took advantage of the extensive driver line collection in *Drosophila* (Luan et al., 2006; Jenett et al., 2012; Dionne et al., 2018). We searched for split-GAL4 lines that only labeled a small proportion of all PNs (Yoshi Aso, unpublished data). Using such drivers, we could sequence a few types of PNs at a time, map those cells onto clusters formed by most PNs, and then use differentially expressed markers among them to decode their identities one-by-one.

*split#28-GAL4* labeled two types of PNs—those that project their dendrites to the DC3 and DA4l glomeruli in developing and adult animals (Figure 2A, B; note that PN types are named after the glomeruli they project their dendrites to). We sequenced those PNs (*split#28*+ PNs hereafter) at 24h APF. We chose this stage because this is when different PN types exhibit the highest transcriptome diversity as hinted by the number of clusters seen in Figure 1F (see following sections for more detailed analysis). To visualize sequenced *split#28*+ PNs, we performed dimensionality reduction using 561 genes identified from most 24h PNs using Iterative Clustering for Identifying Markers (ICIM), a unsupervised machine learning algorithm (Li et al., 2017), followed by embedding in the tSNE space. *Split#28*+ PNs (orange dots) fell into two distinct clusters and intermingled with *GH146*+ PNs (grey dots) (Figure 2C). One cluster mapped to previously decoded DC3 PNs (Li et al., 2017), and the other cluster expressed *zfh2* (Figure 2— figure supplement 1A). We validated that this cluster indeed represents DA4l PNs by visualizing the expression of *zfh2* in PNs utilizing an intersectional strategy by combining *zfh2-GAL4, GH146-Flp*, and *UAS-FRT-STOP-FRT-mCD8-GFP* (hereafter referred to as “intersecting with *GH146-Flp*”) (Figure 2—figure supplement 1B).

**Figure 2.**
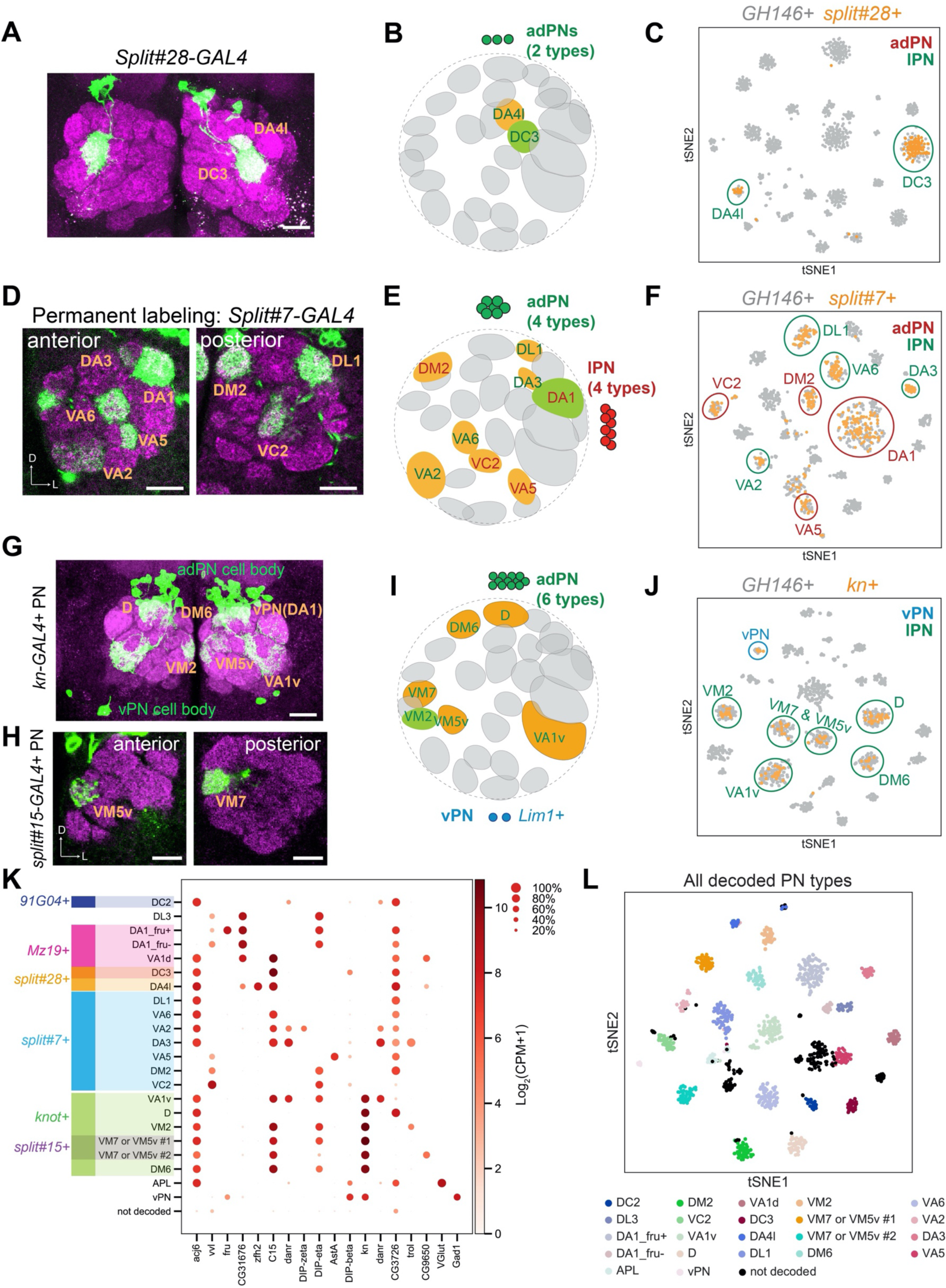
Matching 15 transcriptomic clusters to specific PN types at 24h APF. **(A)** Representative maximum z-projection of confocal stacks of *split#28-GAL4* in adults. Dendrites of *split#28-GAL4*+ PNs target the DC3 and DA4l glomeruli. **(B)** Diagram of *split#28-GAL4*+ PNs. **(C)** tSNE plot showing newly sequenced *split#28-GAL4*+ PNs, which form two clusters that can be assigned to DC3 and DA4l PNs (see also Figure 2–figure supplement 1). **(D)** Representative confocal images of *split#7-GAL4* labeled PNs using permanent labeling strategy. One anterior section and one posterior section of the antennal lobe are shown. Using permanent labeling, we found that this driver is expressed in 8 PN types. Genotype: *split#7-GAL4, UAS-Flp, Actin promoter-FRT-STOP-FRT-GAL4, UAS-mCD8-GFP*. **(E)** Diagram of *split#7-GAL4*+ PNs. *split#7-GAL4* labels 8 types of PNs. 4 from the adPN lineage and 4 from the lPN lineage. **(F)** tSNE plot of *split#7-GAL4* PNs with *GH146*+ PNs (see Figure 2–figure supplement 2 for details on the decoding procedure). **(G)** Representative maximum z-projection of confocal stacks of *kn*+ PNs in the adult. *kn-GAL4* was intersected with *GH146-Flp* to restrict the expression of GAL4 in only PNs. **(H)** Representative confocal images of *split#15-GAL4* in adults, which labels 2 *kn*+ PN types. **(I)** Diagram showing that *kn*+ PNs include 6 types of adPNs (VM2 was decoded) and two vPNs. **(J)** tSNE plot of *kn-GAL4* PNs with *GH146*+ PNs (see Figure 2–figure supplement 3 for details on the decoding procedure). **(K)** Dot plot summarizing drivers and marker genes we used to map 21 transcriptomic clusters to 20 PN types [14 adPNs, 5 lPNs—DA1 PNs form two clusters, one *fru*+ and one *fru–* (Li et al., 2017)—and 1 vPNs] and the anterior paired lateral (APL) neurons at 24h APF. Gene expression level [log2(CPM+1)] is shown by the dot color, and percentages of cells expressing a marker are shown by dot size. **(L)** tSNE plot showing *GH146*+ PNs colored by PN types. Scale bars, 20 μm. Axes, D (dorsal), L (lateral). In panel B, E, and I, orange glomeruli represent PN types of unknown transcriptomic identity prior to this study. Green glomeruli represent PN types whose transcriptomic identity were previously decoded.

*split#7-GAL4* labeled 3 types of PNs in the adult stage (Figure 2—figure supplement 2A). However, when we sequenced cells labeled by this GAL4 line at 24h APF and visualized them using tSNE, 8 distinct clusters were found (Figure 2F). We reasoned that this could be due to loss of driver expression in adult stage for some PN types. To test this hypothesis and reveal PNs that are labeled by this driver transiently during development, we used a permanent labeling strategy to label all cells that express *split#7-GAL4* at any time of development (*split#7*+ PNs hereafter) by combining it with *UAS-mCD8-GFP, Actin promoter-FRT-STOP-FRT-GAL4*, and *UAS-Flp*. Using this strategy, we observed labeling of 8 types of PNs (Figure 2D), consistent with number of clusters we observed by sequencing. Among *split#7*+ PNs, 4 types belong to the adPN lineage (*acj6*+) and the other 4 types belong to the lPN lineage (*vvl*+) (Figure 2E). Only 1 lPN type, DA1 (*CG31676*+), has previously been decoded (Figure 2—figure supplement 2B). We identified differentially expressed genes among *split#7*+ PNs and obtained existing GAL4 lines mimicking their expression. By intersecting those GAL4 lines with *GH146-Flp*, we mapped all 7 previously unknown transcriptomic clusters to 7 PN types (Figure 2—figure supplement 2 C–H; see legends for detailed description).

In addition to screening through collections of existing driver lines, we also utilized scRNA-seq data to find drivers that label a subpopulation of PNs. One such marker we found was the gene *knot* (*kn*), which was expressed in 7 transcriptomic clusters among all *GH146*+ PNs (Figure 2—figure supplement 3A). One of the *kn*+ clusters expressing *trol* has been previously mapped to VM2 PNs (Li et al., 2017). When *kn-GAL4* was intersected with *GH146-Flp*, 6 types of adPNs (*acj6*+) and several vPNs (*Lim1*+) were labeled (Figure 2G, J). Among the 6 adPN types, VM7 and VM5v PNs were also labeled by *split#15-GAL4* (Figure 2H). Although it has been previously reported that *GH146-GAL4* is not expressed in VM5v PNs (Yu et al., 2010), labeling of these PNs when *GH146-Flp* was intersected with either *kn-GAL4* or *split#15-GAL4* indicates that *GH146-Flp* must be expressed in VM5v PNs at some point during development. Using *split#15-GAL4*, we were able to decode the two clusters to be either VM7 or VM5v PNs (Figure 2–figure supplement 3B). Due to the lack of existing GAL4 drivers for differentially expressed genes between these two clusters, we could not further distinguish them so far, but we could create new GAL4 drivers to decode their identities in future studies. Other than these two clusters, we were able to match transcriptomic clusters and glomerular types for the rest of adPNs one-to-one (Figure 2–figure supplement 3C-E). In addition to excitatory PNs, one *kn*+ vPN type innervated DA1 glomerulus (because DA1 glomerulus is innervated only by lPNs and vPNs, not adPNs). We found that *DIP-beta* was expressed in one *kn*+ vPN cluster but not in lPNs innervating DA1 glomerulus (Figure 2—figure supplement 3F, G). Intersecting *DIP-beta-GAL4* with *GH146-Flp* confirmed that *DIP-beta*+ vPN indeed targeted their dendrites to DA1 glomerulus, illustrating the *DIP-beta*+ vPN cluster to be DA1 vPNs (Figure 2—figure supplement 3H).

In summary, by sequencing a small number of known PN types at a time and analyzing the expression pattern of differentially expressed genes, we have now mapped a total of 21 transcriptomic clusters corresponding to anatomically defined PN types at 24h APF (Figure 2K, L). Ultimately, we aimed to match the transcriptomes of the same types of PNs across development. Prior to achieving this goal, we carried out global analysis of gene expression changes across development, which could help us reliably identify transcriptomic clusters representing different PN types at different developmental stages.

### Global gene expression dynamics across four developmental stages

All sequenced PNs segregated into different clusters according to their developmental stages using unbiased, over-dispersed genes for clustering (Figure 1F) regardless of PN types. Even when we used the genes identified by ICIM for clustering, which emphasizes the differences between different PN types (Li et al., 2017), we still observed that individual PNs were separated principally by developmental stages (Figure 3A). Together, these observations illustrate global transcriptome changes of PNs from pupa to adult.

**Figure 3.**
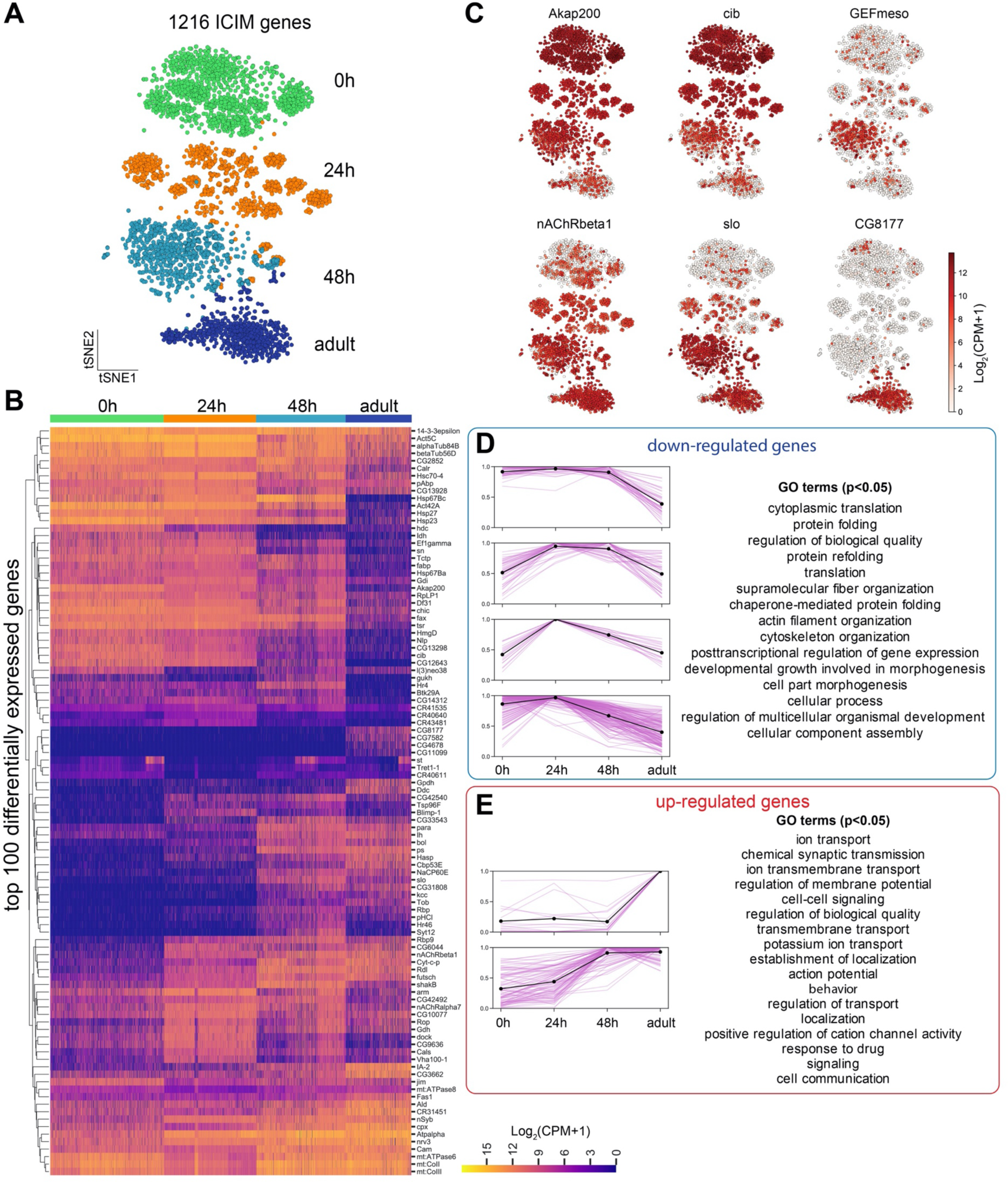
Global-level gene expression dynamics of PNs. **(A)** Visualization of PNs from 4 different developmental stages: 0h APF, 24h APF, 48h AFP, and adult sequenced using either *VT033006-GAL4* or *GH146-GAL4*. tSNE dimensionality reduction was performed using 1216 genes identified by iterative clustering for identifying markers (ICIM) among them. **(B)** Hierarchical heatmap showing the expression of the top 100 out of 474 differentially expressed genes identified among PNs of different developmental stages. **(C)** Examples of the expression of the dynamic genes. Cells are colored according to the expression level of each gene. **(D, E)** Top 474 differentially expressed genes can be divided into 8 groups based on their dynamic profiles–2 groups without obvious developmental trend (not shown), 4 groups of down-regulated genes (D), and 2 groups of up-regulated genes (E). Pink lines represent individual genes and the black line shows mean expression of genes in each group. The highest expression is normalized as 1 for all genes. GO terms for developmentally up-regulated and down-regulated genes are shown on right.

To understand what types of genes drive this separation, we searched for genes that were differentially expressed in different developmental stages (Figure 3B, C). We clustered the genes into different groups based on their expression pattern throughout development. Six groups of genes showed clear developmental trends—four groups were down-regulated from pupa to adult and two groups were up-regulated (Figure 3D–E). Consistent with our previous knowledge, neural development-related genes, including those with functions in morphogenesis and cytoskeleton organization, were enriched in developing PNs; genes related to synaptic transmission, ion transport, and behavior, on the other hand, were up-regulated in mature PNs (Li et al., 2017; Li et al., 2020b).

### Single-cell transcriptomes of PNs reveal dominant biological processes at different stages of development

Because PN transcriptomes exhibited global development-dependent dynamics, we needed to find a method to reliably and consistently classify transcriptomic clusters representing different PN types at all stages. We first identified informative genes for clustering from each stage using ICIM and used them for further dimensionality reduction. However, using this method, we obtained different numbers of clusters at each stage (Figure 4A). Closer examination of each stage revealed unique biological features of PN development.

**Figure 4.**
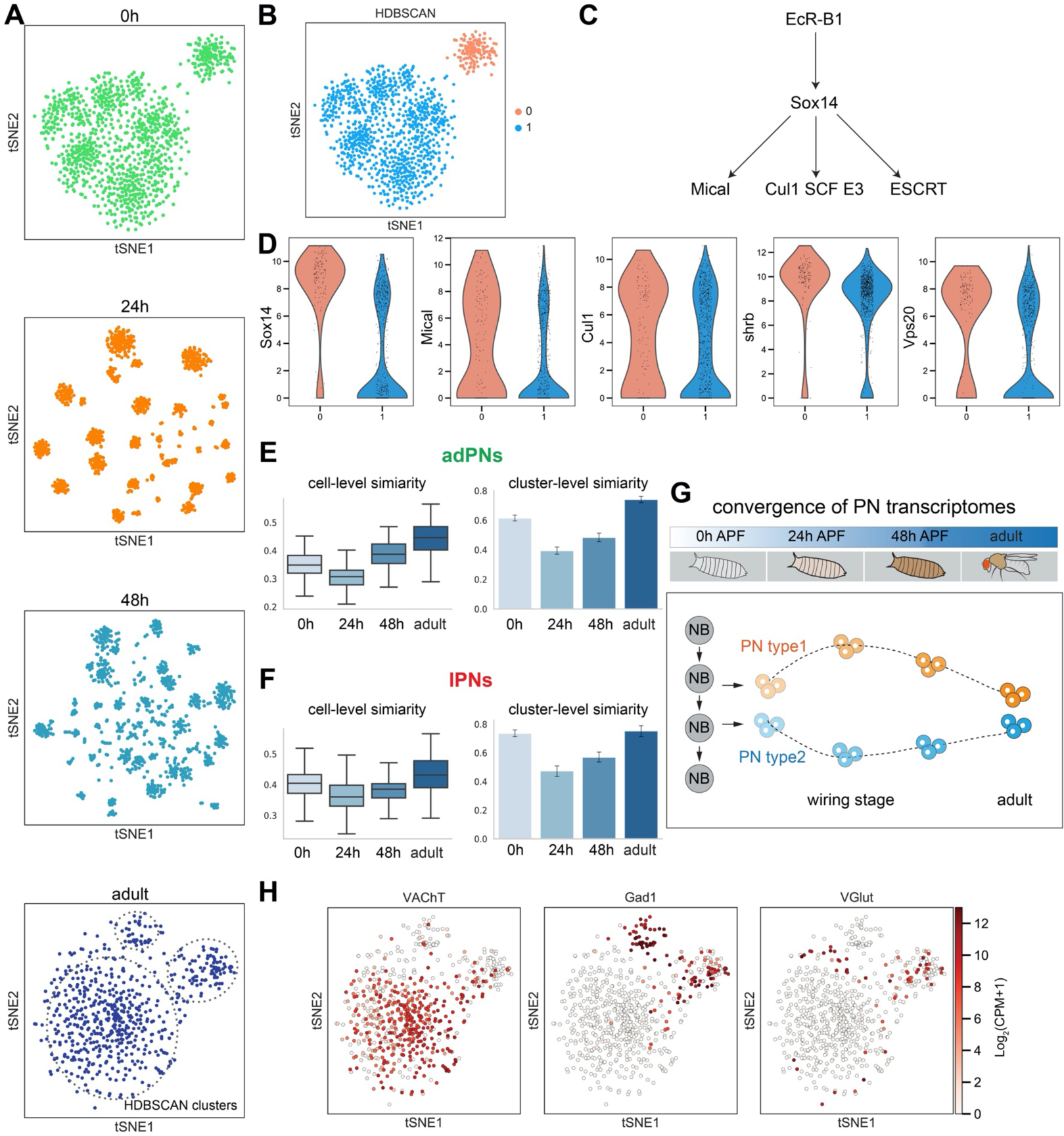
PN transcriptomes show distinct features at different stages of development. **(A)** Visualization of most PNs from 0h APF, 24h APF, 48h APF, and adults using tSNE based on genes identified by ICIM at each stage. Adult clusters (circled) are identified using HDBSCAN. **(B)** Clustering of 0h APF PNs using HDBSCAN identified two clusters. **(C)** Part of the molecular pathways critical for neurite pruning in *Drosophila*. **(D)** Genes whose function have been implicated in neurite pruning have higher expression in cluster 0: *Sox14* (p-value: 5.01E-51), *Mical* (p-value: 1.49E-09), *Cul1* (p-value: 8.15E-4), *shrb* (p-value: 6.37E-19) and *Vps20* (p-value: 1.23E-17) (Mann-Whitney U test). **(E, F)**. PN transcriptomic similarity calculated at the cell level (mean inverse Euclidean distance calculated using 1216 ICIM genes identified from PNs of all 4 stages) and the cluster level (Pearson correlation calculated using differentially expressed genes identified from 24h PN clusters) for adPNs (E) [0h APF: 587 cells, cell-level similarity (mean ± standard deviation): 0.350 ± 0.036, 15 clusters, cluster-level similarity (mean ± standard deviation): 0.615 ± 0.160; 24h APF: 547 cells, cell-level similarity: 0.292 ± 0.041, 15 clusters, cluster-level similarity: 0.395 ± 0.189; 48h APF: 301 cells, cell-level similarity: 0.377 ± 0.046, 13 clusters, cluster-level similarity: 0.484 ± 0.212; adult stage: 209 cells, cell-level similarity: 0.422 ± 0.058, 15 clusters, cluster-level similarity: 0.741 ± 0.129] and lPNs (F) [0h APF: 484 cells, cell-level similarity: 0.402 ± 0.052, 10 clusters, cluster-level similarity: 0.736 ± 0.129; 24h APF: 354 cells, cell-level similarity: 0.360 ± 0.056, 10 clusters, cluster-level similarity: 0.474 ± 0.057; 48h APF: 296 cells, cell-level similarity: 0.385 ± 0.043, 10 clusters, cluster-level similarity: 0.570 ± 0.171; adult stage: 191 cells, cell-level similarity: 0.444 ± 0.057, 8 clusters, cluster-level similarity: 0.754 ± 0.141)] **(G)** Schematic summary of the convergence of PN transcriptomes from early pupal stage to adulthood. PN diversity peaks during circuit assembly around 24h APF and gradually diminishes as they develop into mature neurons. **(H)** Expression of *VAChT, Gad1*, and *VGlut* in adult PNs.

At 0h APF, PNs always formed two distinct clusters—a larger cluster consisting of both adPNs and lPNs, and a smaller one with only adPNs (Figure 4B, Figure 4—supplement 2A). As introduced earlier, although all lPNs and many adPNs are born during the larval stage, some adPNs are born during the embryonic stage. We hypothesized that the smaller cluster could represent embryonically born PNs, which undergo metamorphosis including the pruning of their dendrites and axons (Marin et al., 2005). Neurite pruning in *Drosophila* depends on the function of the steroid hormone ecdysone receptor (EcR) (Levine et al., 1995; Thummel, 1996; Schubiger et al., 1998; Lee et al., 2000) cell autonomously (Lee et al., 2000). Upon binding of the steroid hormone ecdysone, EcR and its co-receptor Ultraspiracle (Usp) form a complex to activate a series of downstream targets, including a transcription factor called Sox14, which in turn promotes expression of the cytoskeletal regulator Mical and Cullin1 SCF E3 ligase (Figure 4C) (Lee et al., 2000; Kirilly et al., 2009; Kirilly et al., 2011; Wong et al., 2013). To test our hypothesis, we examined the expression of genes which are known to participate in neurite pruning and genes that showed elevated expression in the mushroom body γ neurons during pruning (Alyagor et al., 2018). We found that *Sox14, Mical, Cullin1*, and two sorting complexes required for transport (ESCRT) genes—*shrb* and *Vps20*, indeed showed higher expression levels in the smaller cluster (Figure 4D). We also confirmed our hypothesis by mapping two types of embryonically born PNs, DA4l and VA6 PNs, to this smaller cluster (Figure 4—figure supplement 2B; see mapping details in Figure 7).

At 24h APF, we observed the highest number of clusters reflecting different PN types. Moreover, dimensionality reduction using the top 2000 overdispersed genes also showed more distinct clusters at this timepoint compared to the others (Figure 4—figure supplement 1). Quantifications of transcriptomic similarity among PNs at each stage indeed confirmed the highest diversity among PNs at 24h APF (Figure 4E–G). This is likely explained by the fact that at this stage, PNs refine their dendrites to specific regions and begin to prepare themselves as targets for their partner ORN axons. Both processes require high level of molecular diversity among different PN types to ensure precise wiring, warranting more distinction between their transcriptomes at this stage.

In contrast to the high transcriptomic diversity in 24h APF PNs, adult PNs only formed three clusters (Figure 4A bottom, indicated by dashed lines). The three clusters represent excitatory PNs (marked by *VAChT*), and two *Gad1*+ GABAergic inhibitory cell types—vPNs and APL neurons (*VGlut*+), respectively (Figure 4H). This is likely because after wiring specificity is achieved, all excitatory PNs may perform similar functions in comparison with the other two neuronal types.

Thus, at three different developmental stages, the differentially expressed genes we identified all revealed the most defining biological processes those neurons are undertaking. Our observations showed that PN transcriptomes reflect the pruning process of embryonically born PNs at 0h APF, PN type and wiring distinction at 24h APF, and neurotransmitter type in adults.

### Identifying PN types at all developmental stages

With the exception of the 24h APF PNs, gene sets identified from each of the other stages could not resolve distinct clusters reflecting PN type diversity (Figure 4). Therefore, we tried to use the genes identified by ICIM from 24h APF PNs to cluster PNs of the other stages. We found that this gene set outperformed all other gene sets in separating different PN types at all timepoints (Figure 5A). In fact, most gene sets found by different methods at 24h APF, including overdispersed genes, ICIM genes, as well as differentially expressed genes between different clusters, exceeded gene sets identified at other stages for clustering PNs according to their types (data not shown), further confirming that transcriptomes of 24h APF PNs carry the most information for distinguishing different PN types, even for other developmental stages.

**Figure 5.**
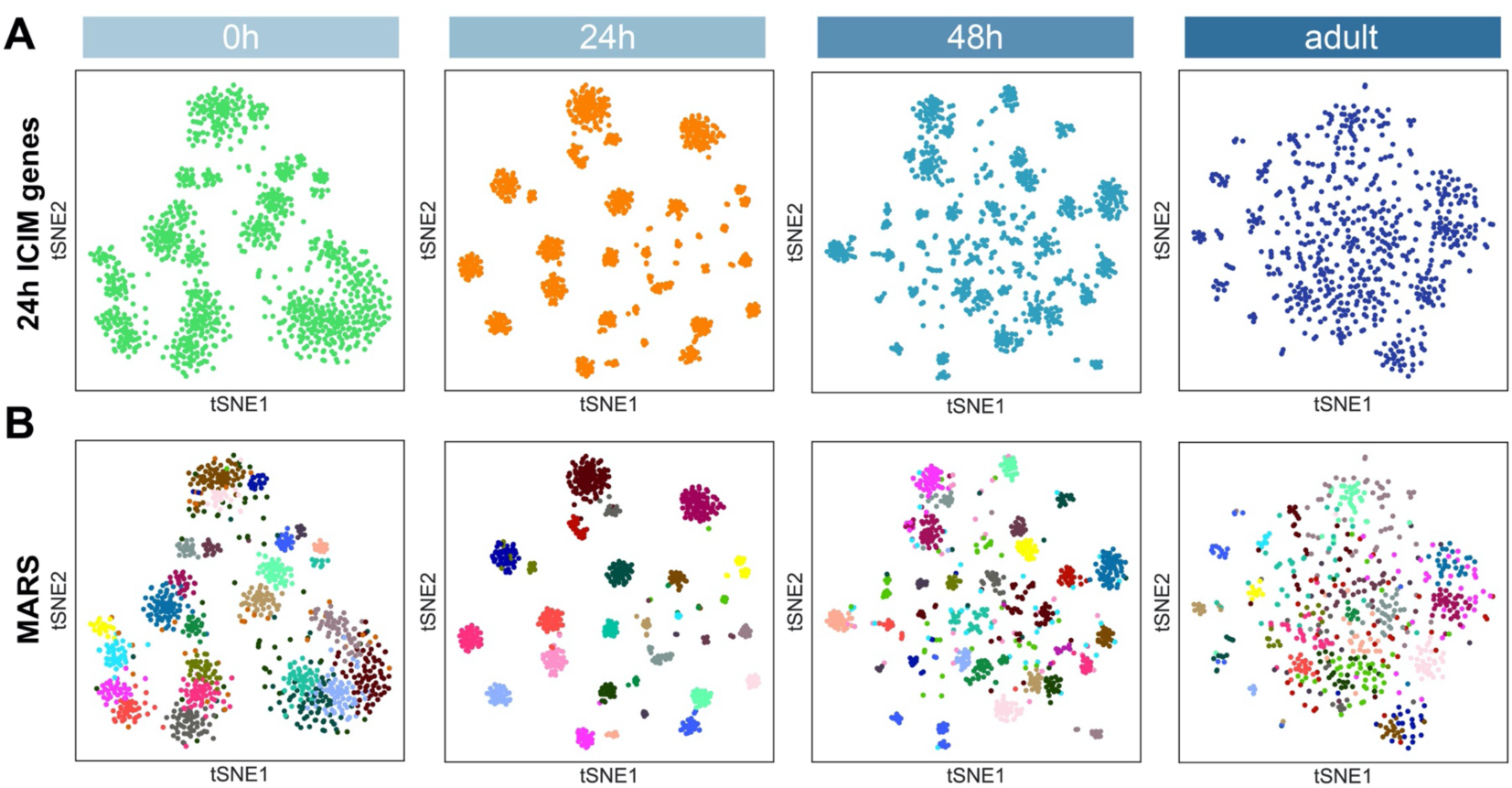
PN type identification by MARS. **(A)** Dimensionality reduction of most PNs at 4 developmental stages by 561 ICIM genes found at 24h APF. **(B)** PN types identified by MARS. Different PN types are illustrated in different colors.

Following this observation, we decided to use differentially expressed genes between 24h PN clusters for PN-type identification for all stages. We applied <u>m</u>et<u>a</u>-learned representations for <u>s</u>ingle cell data (MARS) for identifying and annotating cell types (Brbić et al., 2020). MARS learns to project cells using deep neural networks in the latent low-dimensional space in which cells group according to their cell types. Using this approach, we found ∼30 cell types in each stage (Figure 5B). Independently, we also validated MARS cluster annotations using two distinct methods: HDBSCAN clustering based on tSNEs or Leiden clustering based on neighborhood graphs (Figure 5—figure supplement 1) (Blondel et al., 2008; Levine et al., 2015; Traag et al., 2019). Clusters identified by HDBSCAN and Leiden largely agreed with MARS annotations, confirming the reliability of MARS. We compared cluster annotations by these three methods to known PN types at 24h APF (Figure 5–figure supplement 1C) and found that MARS performed better at segregating closely related clusters representing multiple PN types (Figure 5–figure supplement 1D), demonstrating the robustness of MARS at identifying unique cell types.

### Matching the same PN types across four developmental stages

We next sought to match transcriptomes of the same PN type across different developmental stages. To develop reliable approaches to perform this task, we first used *kn*+ PNs as test case. We collected PNs labeled by *kn-GAL4* from 24h APF, 48h APF, and adult brains for scRNA-seq (Figure 6A). Dimensionality reduction of these cells showed a consistent number of clusters across stages (Figure 6B). One exception is an extra vPN cluster observed at 48h APF and adult stages. This discrepancy with 24h APF data is likely caused by the lower number of vPNs sequenced at 24h APF.

**Figure 6.**
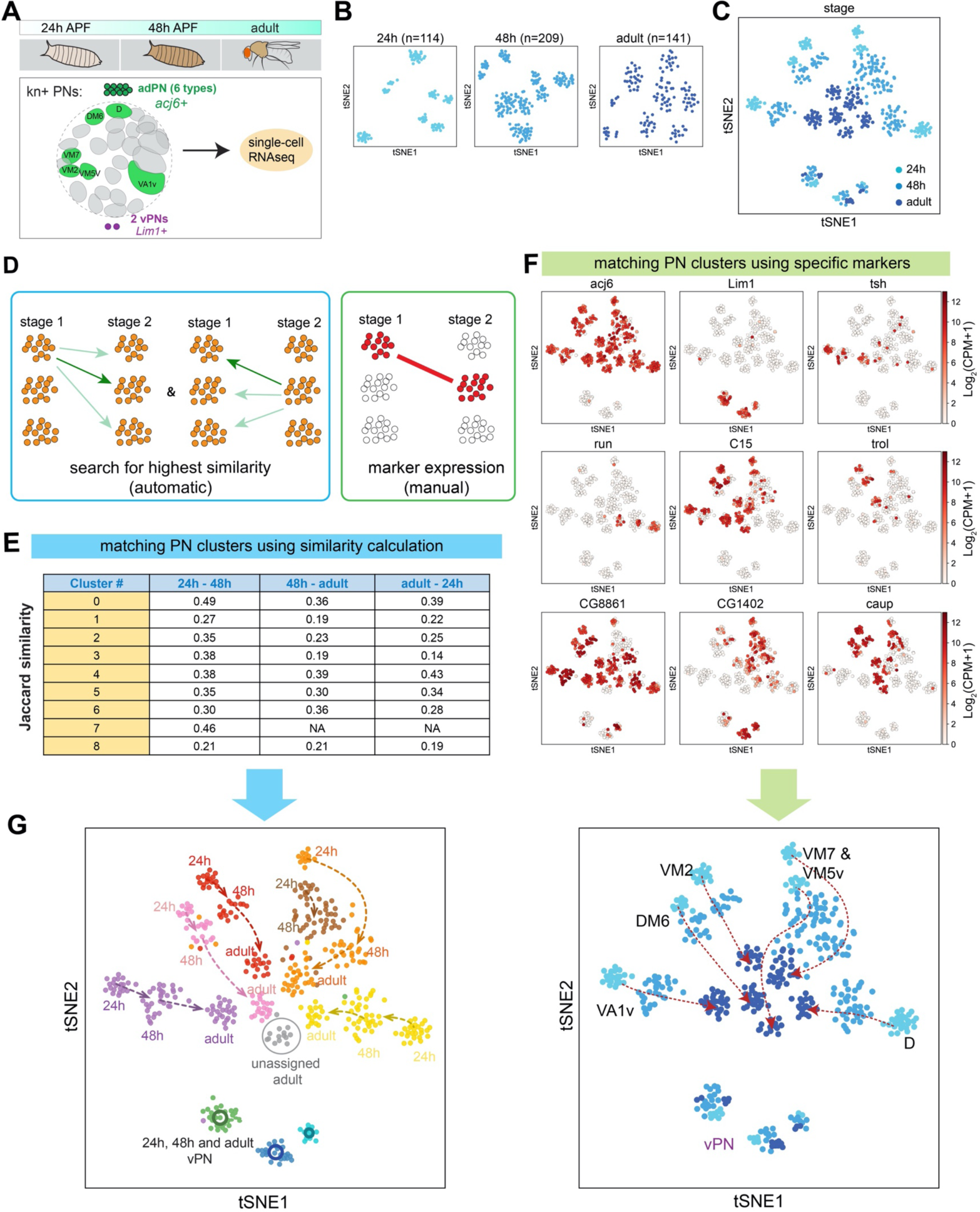
Two complementary approaches to match transcriptomic clusters representing same PN types at different developmental stages. **(A)** scRNA-seq was performed for *kn*+ PNs from 3 different developmental stages: 24h APF, 48h APF, and adult. **(B)** tSNE plots showing *kn*+ PNs from three different stages, plotted separately. Cells are clustered according to 24h ICIM genes. Cell numbers are indicated. **(C)** *kn*+ PNs from three different stages plotted in the same tSNE plot. Cells are clustered according to 24h ICIM genes. **(D)** Two approaches were used for matching the same PN types at different stages: 1) automatic prediction by calculating the transcriptomic similarity between clusters at two stages 2) manual matching of clusters using specific markers or marker combinations. **(E)** Jaccard similarity index of automatically matched transcriptomic clusters from different stages. **(F)** Examples of markers used to manually match transcriptomic clusters representing the same PN types across different stages. **(G)** All *kn*+ PN types (6 adPNs and 3 vPNs) are matched from three different stages. Two independent approaches (automatic and manual) produced similar results.

When *kn*+ PNs from all three stages were plotted together, all adPNs (*acj6*+ clusters on the upper side) formed relatively distinct clusters and did not intermingle with adPNs from the other timepoints (Figure 6C), reflecting substantial changes in the transcriptome of the same type of PNs across development. To match the same type of PNs, we took two independent approaches (Figure 6D). In the first approach, clusters were automatically matched based on their transcriptomic similarity. Briefly, we identified a set of genes that were differentially expressed in each cluster compared to all the rest at the same stage. Then, we calculated the percentage of genes shared between each pair of clusters across two stages (Jaccard similarity index) (Figure 6E). If two clusters from two stages both had the highest similarity score with each other, we considered them to be matched. In the second approach, we used markers that were expressed in a consistent number of clusters at each stage. Those markers, or marker combinations, were used to manually match the same type of PNs (some example markers used are shown in Figure 6F). Using these two approaches, we were able to match the same types of PNs across three developmental stages, and the results from the two approaches consistently agreed with each other (Figure 6G). In addition, these data further validated an earlier conclusion (Figure 4) that as development proceeds from 24h APF and 48h APF to adults, the transcriptomic difference between identified PN types becomes smaller (Figure 6G; quantified in Figure 6—figure supplement 1).

We next applied the same approaches for matching *kn*+ PN types across 3 stages to match most PNs (sequenced using either *GH146-GAL4* or *VT033006-GAL4*) across 4 stages (Figure 7A). In addition to marker gene expression, we also used subset of PNs we had sequenced from different stages to manually match PN types (Figure 7—figure supplements 1A–D). For the manually matched PN types with known identity, we summarized markers and marker combinations we used in a dot plot, where both average expression as well as percentage of cells expressing each marker were shown (Figure 7–figure supplement 2). Using both manual and automatic approaches, we were able to match many PN types across 2 or more developmental stages (Figure 7B), which includes 18 PN types that we have decoded in Figure 2 and 7 transcriptomic clusters with unknown identity. The majority of the PNs we matched were confirmed mutually by both the automatic (transcriptomic similarity-based) and manual (marker-based) methods (Figure 7C and Figure 7– figure supplement 1E).

**Figure 7.**
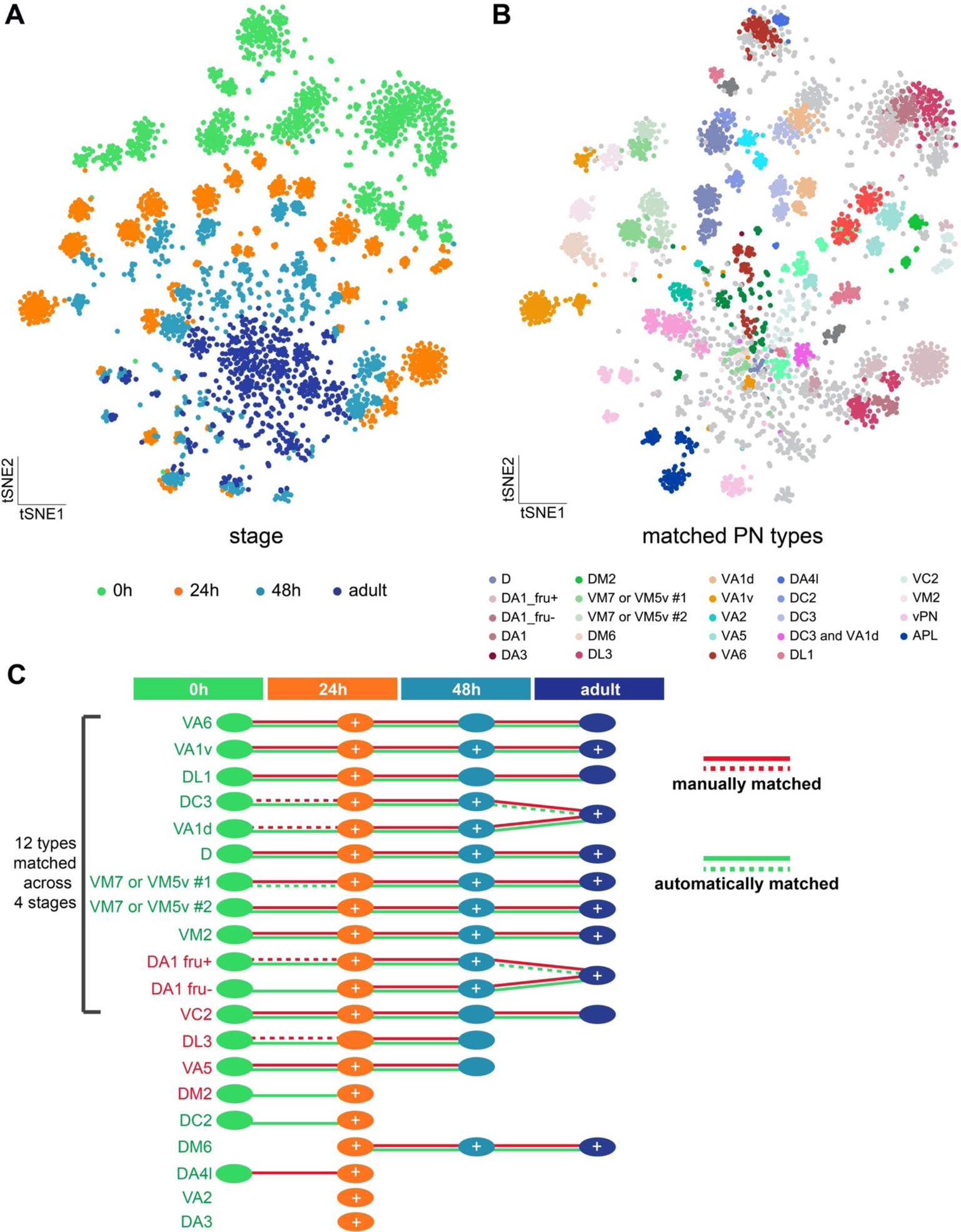
Matching transcriptomic cluster representing the same PN types across four developmental stages. **(A)** Visualization of most PNs in 4 different developmental stages: 0h APF, 24h APF, 48h APF, and adult. 561 ICIM genes at 24h APF PNs are used for dimensionality reduction. **(B)** Visualization of the same types of PNs at all developmental stages. Clusters with the same color represent same neuronal type. Light grey dots indicate cells that have neither been decoded nor matched. **(C)** Summary of transcriptomic clusters mapped to known PN types at different developmental stages. Solid red-lines indicate clusters we can unambiguously match using marker combinations; dashed red-lines indicate PN types we can narrow down to less than 3 transcriptomic clusters. Solid green-lines indicate clusters that are two-way matched automatically (two clusters from two stages are the most similar to each other); dashed green-lines indicates clusters that are one-way matched automatically (one cluster is the most similar with the other but not the other way around). Circles with white “+” on it indicate this PN type have been sequenced and confirmed at that stage using additional GAL4 lines (see figure 7—figure supplement 1).

### Gene expression dynamics in a type-specific manner

Matching the same PN types across multiple developmental stages enabled us to investigate gene dynamics in each PN type. Genes with temporal dynamics in PNs on the bulk level displayed features of neurite growth during development and synaptic transmission in adult stage (Figure 3). However, not many genes known to be involved in wiring-specificity were observed in the differentially expressed gene list when we only considered developmental stage (but not PN type) as a variable. We hypothesized that genes with wiring function might display type-specific dynamics that could not be observed on the global level. Thus, we sought to systematically identify those genes.

We first focused on 6 types of *kn*+ adPNs from 3 stages. We searched for two categories of type-specific dynamic genes: (i) dynamic-dynamic genes, and (ii) dynamic-stable genes. We defined dynamic-dynamic genes to be those that show significant changes in the opposite directions between at least two PN types at two stages, and dynamic-stable genes to be those that have altered expression level in some PN types but maintain stable expression or are not expressed in all stages (Figure 8A). We identified 26 dynamic-dynamic genes and 50 dynamic-stable genes with false discovery rate (FDR) < 0.01 among *kn*+ PNs (Figure 8B, C). Two examples of these type-specific dynamic genes*—Pvf3*, a ligand for the receptor tyrosine kinase encoded by *PvR*, and *rad*, a Rap-like GTPase activating protein—are shown in Figure 8D. The expression of *Pvf3* peaked at different timepoints for D PNs (at 0h APF), VA1v PNs (48h APF), and VM7 or VM5v PNs (in adults). The expression of *rad* decreased in VA1v PNs and increased in VM2 PNs from 48h APF to the adult stage. Interestingly, more than half of the dynamic-dynamic genes (14 out of 26) are cell surface molecules (CSMs) and transcription factors (TFs). Consistent with our hypothesis, both CSMs and TFs are known to play critical roles in PN wiring (Hong & Luo, 2014; Li et al., 2017).

**Figure 8.**
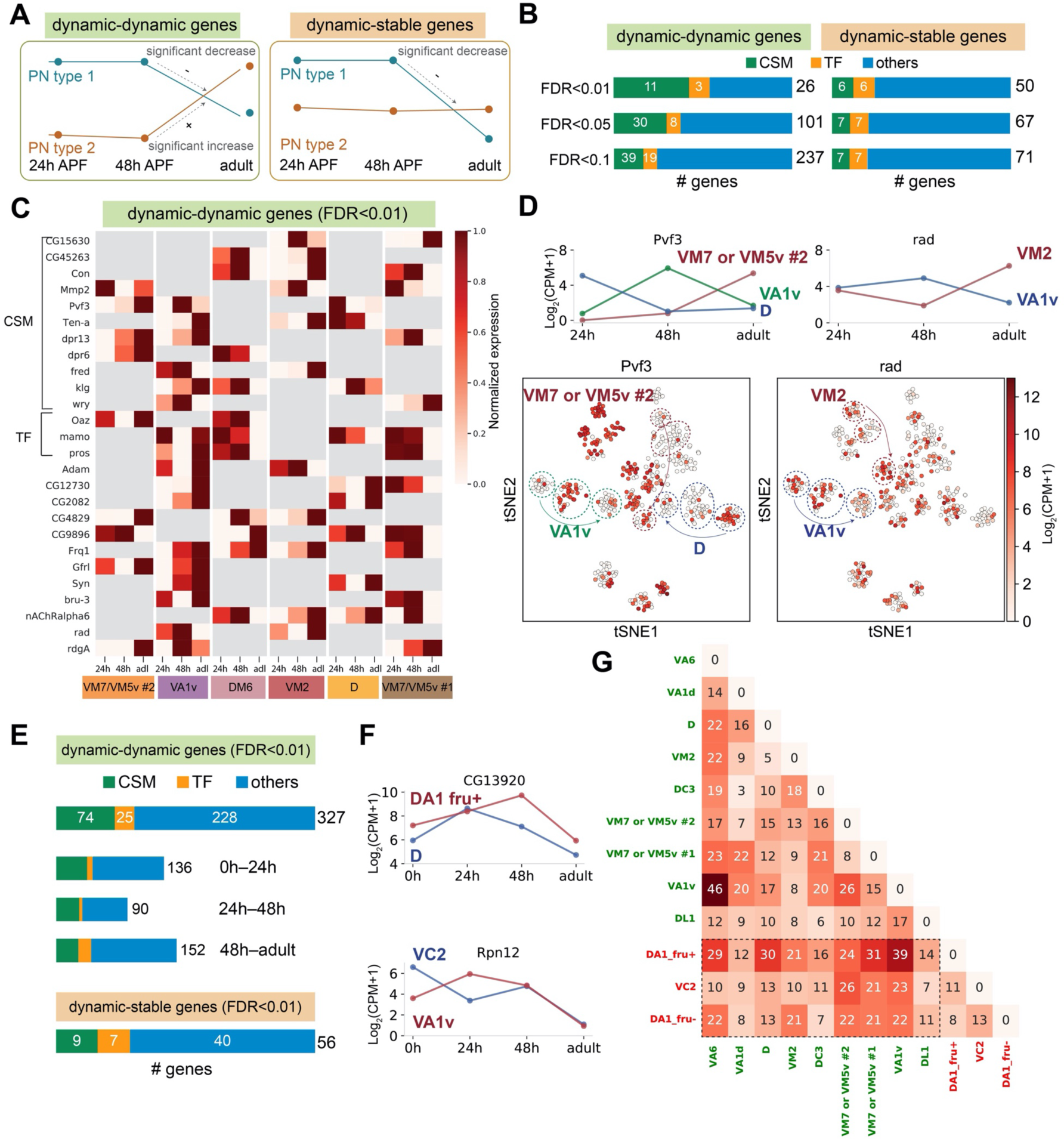
Type-specific dynamic genes among PNs. **(A)** Illustration of an example of dynamic-dynamic genes (left) and dynamic-stable genes (right). **(B)** Number of genes with type-specific dynamics found in *kn*+ adPNs using different false discovery rate (FDR) cutoffs. We highlighted the number of cell surface molecules (CSMs) or transcription factors (TFs). **(C)** Heatmap of dynamic-dynamic genes found among *kn*+ adPNs with FDR < 0.01. Each row shows expression patterns of a gene in different PN types from 24h APF to the adult stage. Gray color means no significant change were observed in that PN type across development. The highest expression is normalized to 1. **(D)** Examples of dynamic-dynamic genes found among *kn*+ PNs. Top: average expression of *Pvf3* and *rad* in PN types with different dynamics. Bottom: tSNE plot of *kn*+ PNs colored by the expression level of *Pvf3* and *rad*. **(E)** Number of genes with type-specific dynamics found among the PN types we matched in all four stages (FDR < 0.01). For the 327 dynamic-dynamic genes, we also categorized them according to the two-stage transitions the PN-type specific dynamics are observed (note that some genes have type-specific dynamics in more than one transition). **(F)** Examples of dynamic-dynamic genes reported in (E). **(G)** Number of dynamic-dynamic genes found between each pair of the 12 decoded PN types. Names of adPNs are in green and names of lPNs are in red. Comparison between adPNs are lPNs are highlighted in the dashed box.

Next, we extended this analysis to more PN types. We focused on 13 PN types that were matched across all 4 developmental stages (12 PN types with known identity and 1 with unknown identity). The increased number of PN types and the additional timepoint produced more type-specific dynamic genes. In particular, at FDR < 0.01 we identified 327 dynamic-dynamic genes (Figure 8E–F). Among the 327 dynamic-dynamic genes, we compared the gene distribution at 3 transitions: 0h to 24h, 24h to 48h, and 48h to adult. We found more dynamic genes during the first and last transitions compared to the middle one. This is consistent with our expectations because PNs from 0h to 24h, or from 48h to adult, are transitioning into or out of circuit assembly, respectively. We further compared the number of dynamic genes found at all stages between each pair of 12 decoded PN types (Figure 8G). We found that PN types from two different lineages (rectangle-bound corner) tended to have more dynamic genes between each other than PN types within the same lineage. However, there were exceptions—for example, VA6 and VA1v PNs are both from the adPN lineage but possessed the highest number of type-specific dynamic genes. This is likely because VA6 and VA1v PNs are born during different developmental stages (born in embryos vs larvae, respectively), with VA6 but not VA1v PNs undergoing dendrite and axon pruning followed by re-extension during morphogenesis.

### PN types with adjacent birth order share more similar transcriptomes at early stages of development

Previous works have shown that the PN glomerular types are prespecified by the neuroblast lineages and birth order within each lineage (Jefferis et al., 2001; Yu et al., 2010; Lin et al., 2012) (Figure 9A). Decoding the transcriptomic identities of different PN types at different timepoints allowed us to ask: to what extent is transcriptomic similarity contributed by lineage, birth order, and/or spatial position of their glomeruli? Do these contributions persist through development?

**Figure 9.**
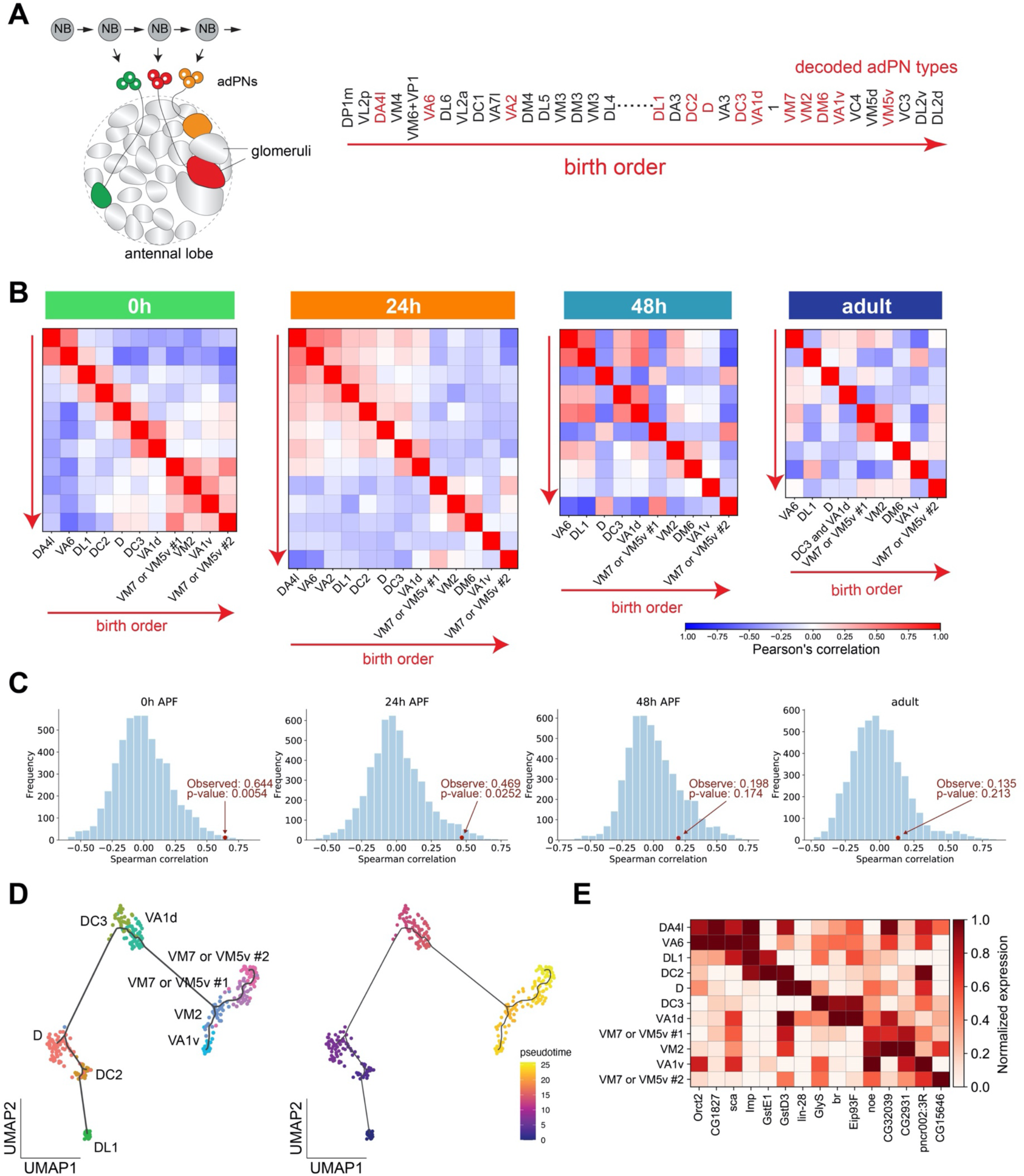
PN types with adjacent birth order share more similar transcriptomes at early pupal stages. **(A)** Different PN types born from a common neuroblast follow a stereotyped sequence. The birth order of PNs determines to which glomerulus their dendrites target. The birth order of adPNs are shown on right. PN types with known transcriptomic identities are highlighted in red. **(B)** Correlation matrix of the transcriptomes of adPNs with known identities (Pearson’s correlation). PN types are ordered according to their birth order. At 0h and 24h APF, PN types with birth orders adjacent to each other exhibit the highest correlations in their transcriptomes, as indicated by high correlations in boxes just off the diagonal line. **(C)** Results of permutation test under the null hypothesis that the ranks of adPN transcriptomic similarity do no covary with the ranks of birth order. Observed values is the average Spearman correlation of 8 adPN types decoded in all 4 stages (red dot). The distribution is the average Spearman correlations obtained by randomly permutating the birth order for 5000 iterations (histogram). **(D)** Developmental trajectory analysis showing an unbiased pseudo time of 0h APF adPNs (embryonically born types excluded). The pseudo time roughly matches their birth order. **(E)** Expression levels of 15 genes in adPNs with known identity at 0h APF. These genes have been shown to have temporal expression gradient in PN neuroblasts (Liu et al. 2015). The highest expression is normalized as 1 for all genes.

To address these questions, we performed hierarchical clustering on all excitatory PN clusters we identified from each timepoint. We plotted the dendrogram and the correlation between each pair of clusters (Figure 9—figure supplement 1). We observed some lineage-related similarity between PN types at 0h APF: transcriptomes of PNs from the same lineage tended to be clustered together in the dendrogram and their correlations are higher, although the relationship was not absolute. Such similarity was gradually lost as development proceeded (as inferred by both the dendrogram as well as correlation between PNs from the same lineage). Interestingly, we noticed that some PNs with adjacent birth order appeared to be neighbors in the dendrogram at 0h and 24h APF.

To further investigate the relationship between birth order of PNs and their transcriptomic similarity, we selected all decoded PNs from the anterodorsal lineage, ordered them according to their birth order, and computed their correlation (Figure 9B). 0h APF adPNs showed high correlation between their birth order and their transcriptomic similarity, as indicated by the high correlations in boxes just off the diagonal line. To test if the transcriptomic similarity of adPNs indeed covaries with their birth order, we performed permutation tests, comparing the Spearman correlations between birth-order ranking and transcriptomic similarity ranking (Figure 9C, see Materials and Methods for details). The results confirmed that 0h and 24h APF PNs, but not 48h APF and adult PNs, exhibited high correlations between their birth orders and transcriptomic similarities. In addition, developmental trajectory analysis of adPNs born at the larval stage using Monocle 3 also showed that the unbiased pseudo time recapitulated their birth order (Figure 9D) (Cao et al., 2019).

A previous study profiled the transcriptomes of PN neuroblasts at various larval stages and identified 63 genes with temporal gradients (Liu et al., 2015). Among those genes, the authors have validated that two RNA-binding proteins, Imp and Syp, regulate the fate of PNs born at different times. Therefore, we analyzed expression of these genes at 0h APF to see if any of these genes with temporal gradients has persisted expression in postmitotic PNs. We found 15 out of the 63 genes (including *Imp* but not *Syp*) maintained the same temporal gradient patterns according to their birth order at 0h APF (Figure 9E) but not at the later stages (data not shown). This result suggested that the expression of some birth order-related molecular features, including some cell-fate regulators, were maintained till early pupal stage.

In summary, our data demonstrated that PN types with adjacent birth order shared more similar transcriptomes, illustrating sequential transition of gene expression profiles in PN neuroblasts. Such transcriptomic similarity was maintained at early pupal stages and was gradually lost as PNs mature.

### Differentially expressed genes in different PN types in adults

Our analyses have shown that transcriptomic differences between different PN types diminish as development proceeds (Figure 4). However, different PN types in adults still exhibited some degree of differential gene expression, as demonstrated by the clustering of adult PNs (Figure 5) and the negative correlations observed between some PN types (Figure 9–figure supplement 1D). Such differential expression could be contributed by residual developmentally differentially expressed genes, by new categories of differentially expressed genes in mature PNs reflecting functional differences between different PN types, or a combination of both. To distinguish between these possibilities, we compared differentially expressed (DE) genes among different transcriptomic clusters of PNs at 24h APF and at the adult stage.

About a third of the DE genes were shared between these two stages (Figure 10A). Gene ontology analysis revealed that these shared genes were predominately related to neural development (Figure 10B, middle). In addition, CSMs and TFs were enriched in 24h APF and adult DE genes compared to the entire genome, albeit to a lesser extent for TFs (Figure 10C). These data suggested that some DE genes found among adult PN types were residual developmentally differentially expressed genes.

**Figure 10.**
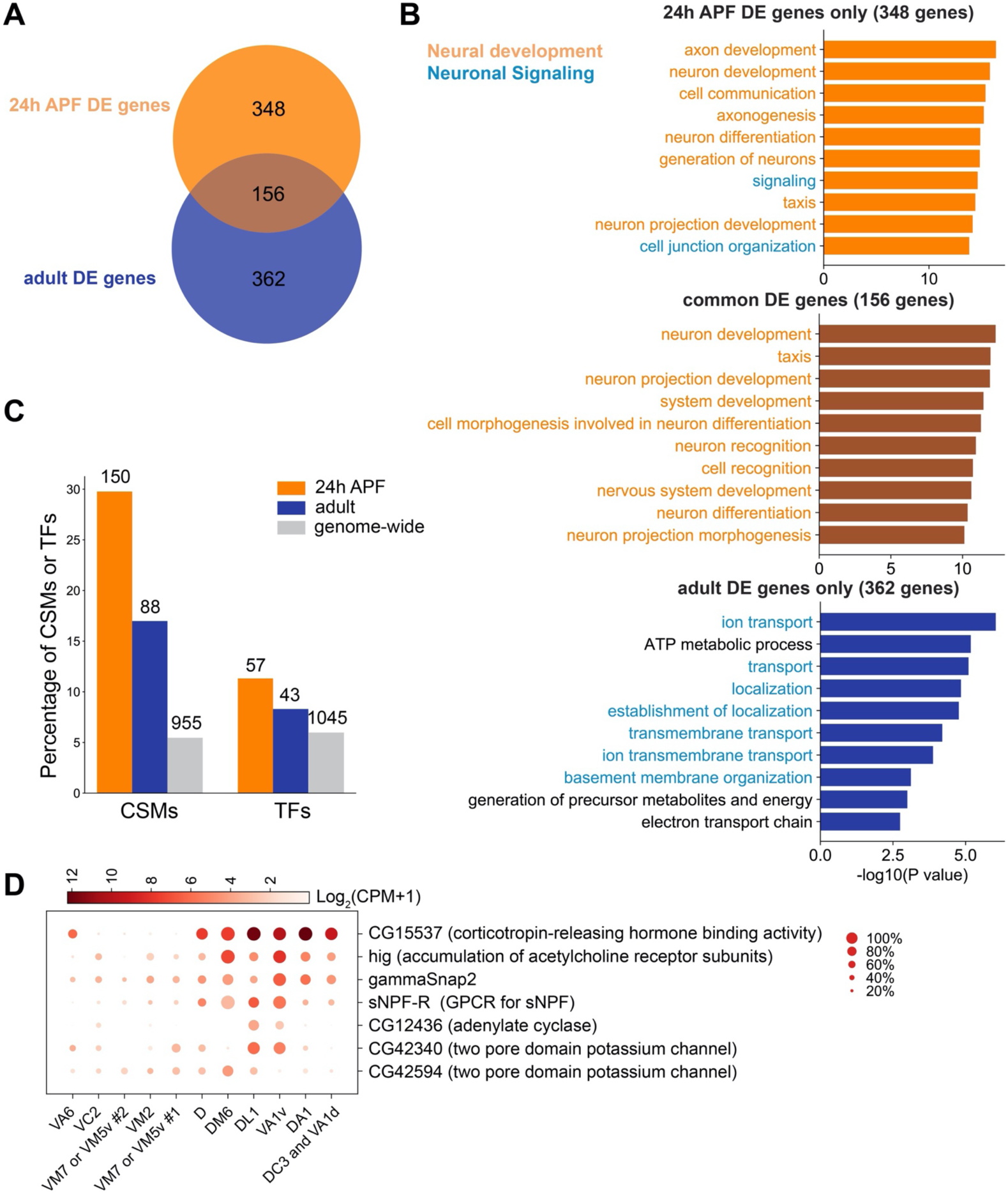
Differentially expressed genes among different PN types in the adult stage. **(A)** Venn diagram of differentially expressed genes (DE genes) at 24h APF (504 genes) and in adults (518 genes). DE genes are genes with adjusted p-value less than 0.01 by Mann-Whitney U test in at least one cluster compared to the rest. **(B)** Top 10 biological process terms of DE genes found in 24h APF PNs only (top), in both 24h APF and adults PNs (middle), and in adult PNs only (bottom). **(C)** Percentage of CSMs or TFs in 24h APF DE genes, adult DE genes, and all *Drosophila* genes. Total numbers of genes within each category are labeled above the bars. 51/88 CSMs and 29/43 TFs of adult DE genes were also found among 24h APF DE genes. **(D)** Dot plot showing the expression of 7 example genes related to neuronal signaling in adult PNs. Example genes were manually selected based on their differential expression pattern among decoded transcriptomic clusters and functions.

Interestingly, many gene ontology terms related to the physiological properties of PNs among the adult only DE genes (Figure 10B, bottom). These include ion channels, G-protein-coupled receptors, and regulators of synaptic transmission (some selected examples are shown in Figure 10D). These results suggested different PN types in adults might exhibit different physiological properties. Future studies can address whether such differences in the adult PN transcriptomes have an impact on their physiological properties.

## Discussion

### Deciphering single-cell transcriptomes for connectivity-defined neuronal types

Traditionally, neurons are classified based on their morphology, physiology, connectivity, and signature molecular markers. More recently, scRNA-seq has allowed classification of cell types based entirely on their transcriptomes. Many studies have illustrated that cell-type classification based on the single-cell transcriptomes largely agrees with classifications by some of the more traditional criteria (Zeng & Sanes, 2017).

For *Drosophila* olfactory PNs, the most prominent type-specific feature is their pre- and post-synaptic connections, which determines their olfactory response profiles and the higher order neurons they relay olfactory information to. Thus, different PN types are largely defined by their differences in their connectivity. We have previously observed that the transcriptomic identity of PNs corresponds well with their types during development, and for three identified PN types, transcriptomic differences peak during the circuit assembly stage (Li et al., 2017). Here, we generalized these findings across many more PN types by showing that transcriptomic differences are the highest around 24h APF, a stage when PNs are making wiring decisions and preparing cues for subsequent ORN-PN matching (Figure 4), and by demonstrating that clustering of PNs according to their types from all stages are best done using differentially expressed genes at 24h APF (Figure 5). Additionally, our data indicate that at certain stages, differences among those type-specific genes can be masked by other genes belonging to pathways of a more dominating biological process (such as neurite pruning at 0h APF for PNs). As a consequence, it may be challenging to identify genes carrying type-specific information at certain timepoints even when sophisticated algorithms are applied, which can lead to underestimation of cell type diversity. Thus, to determine single-cell transcriptomes of connectivity-defined neuronal types such as fly olfactory PNs, it may be a general strategy to first obtain their single-cell transcriptomes during their circuit assembly and then use this information to supervise cell-type classification in other developmental stages, including adults.

### Tracing the same cell type in different states

Both cell types and their biological states can split single-cell transcriptomes into distinct clusters (Zeng & Sanes, 2017; Cembrowski & Menon, 2018; Tasic, 2018). We observed that the same types of PNs of different developmental stages—reflecting different states—indeed exhibit very distinct transcriptomic profiles (Figures 5 and 6). To identify transcriptomic clusters corresponding to the same PN types across multiple timepoints, we developed and applied two complementary methods—one manual based on the marker gene expression, and one automatic based on the similarity between transcriptomic clusters. By applying both methods, we can confidently track the transcriptomes of the same cell type throughout development and study the unique molecular features of each stage.

Our methods can be applied to other single-cell studies where diverse cell types and multiple states are involved. This can be especially useful for tissues with high cellular diversity but lack unique markers for each cell type.

### Using single-cell RNAseq data to identify new candidate molecules for future studies

In this study, we have obtained high-quality single-cell transcriptomes of most excitatory PNs from early pupal stage to adulthood (Figure 1). We have used combinations of markers and drivers to decode the transcriptomic identity of 21 transcriptomic clusters at 24h APF (Figure 2), and matched clusters representing the same PN type across four developmental stages (Figure 7).

Using this rich and well-annotated dataset, researchers can now explore different aspects of PN development and function to identify candidate molecules for future studies. For example, one can search for novel molecules involved in neurite pruning among the differentially expressed genes between the embryonically-born and larval-born PNs at 0h APF (Figure 4B–D). Developmentally enriched genes and genes with type-specific dynamics, on the other hand, can be good candidates for studies on neural development and wiring specificity (Figure 3 and 8). Differentially expressed neuronal signaling genes in adult PNs can be used to explore differences in physiological properties and information processing (Figure 10). In addition, driver lines for specific types of PNs can be made using genes that show consistent expression pattern across different stages (Figure 7–figure supplement 2) to label and genetically manipulate specific PN types. Together with a companion paper on single-cell transcriptomes of olfactory receptor neurons across multiple stages (McLaughlin et al.), these studies have established foundations of gene expression for the two principal types of neurons in the *Drosophila* olfactory system and should catalyze new biological discoveries.

## Methods and Materials

### Key Resource Table

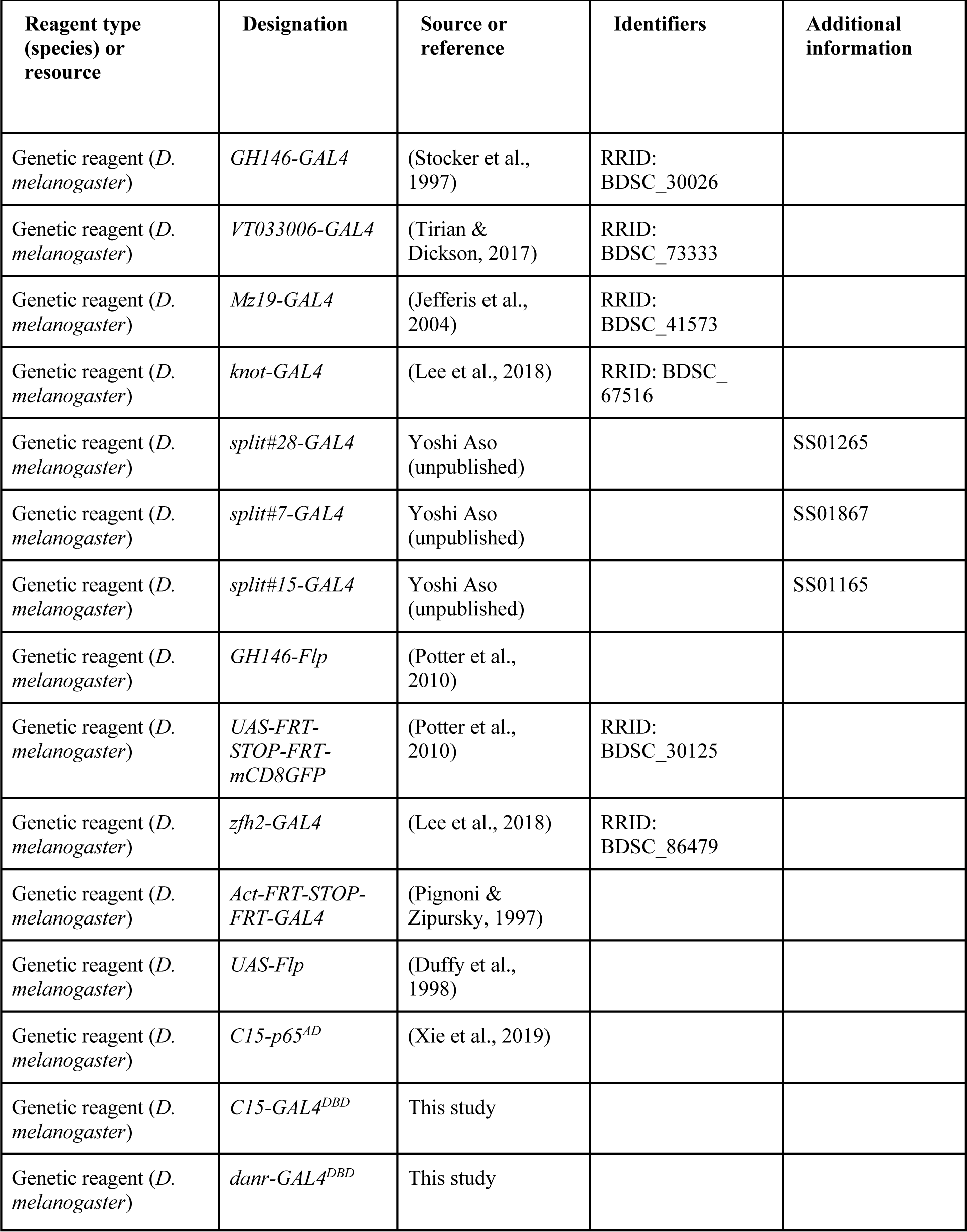
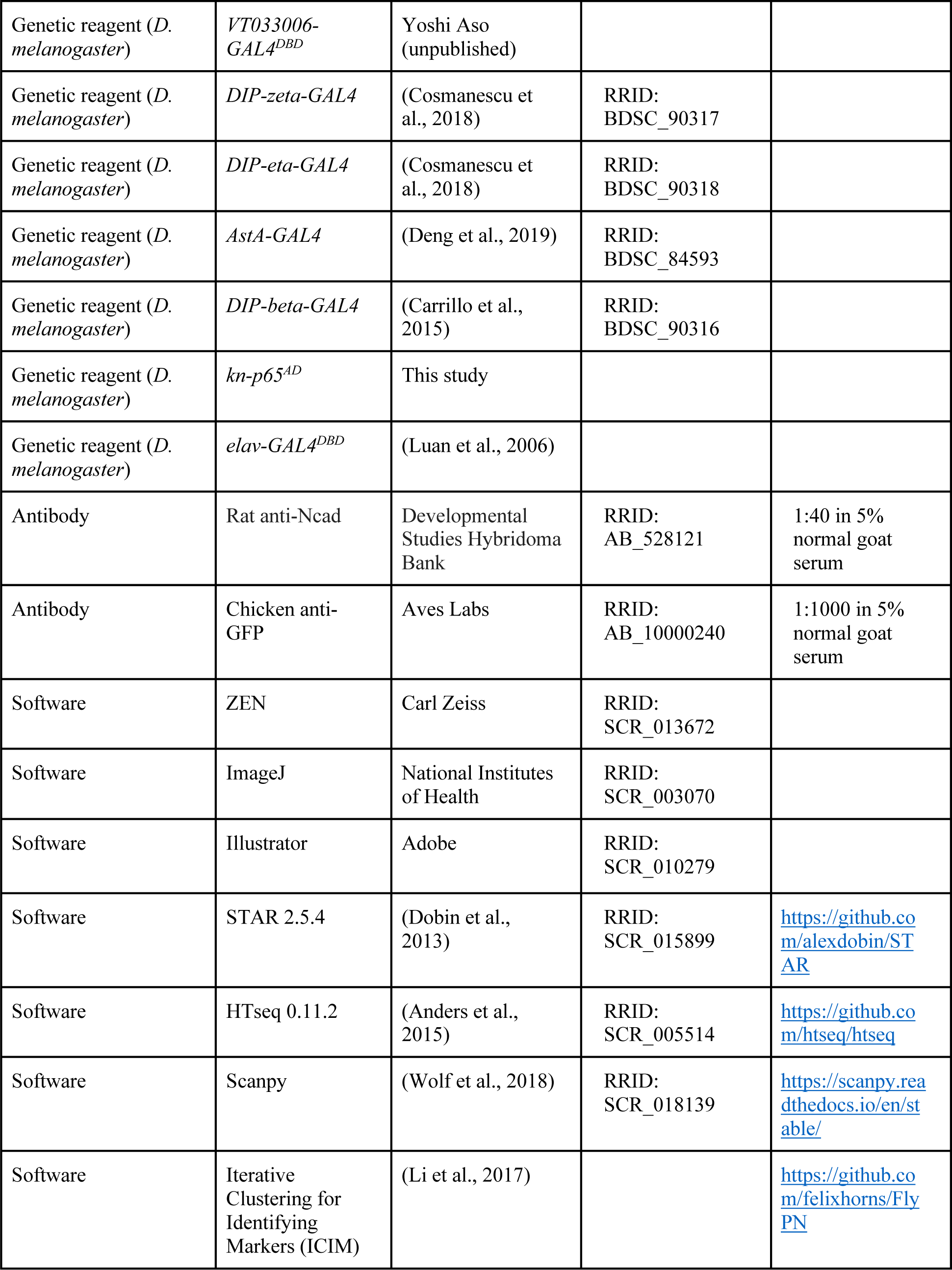
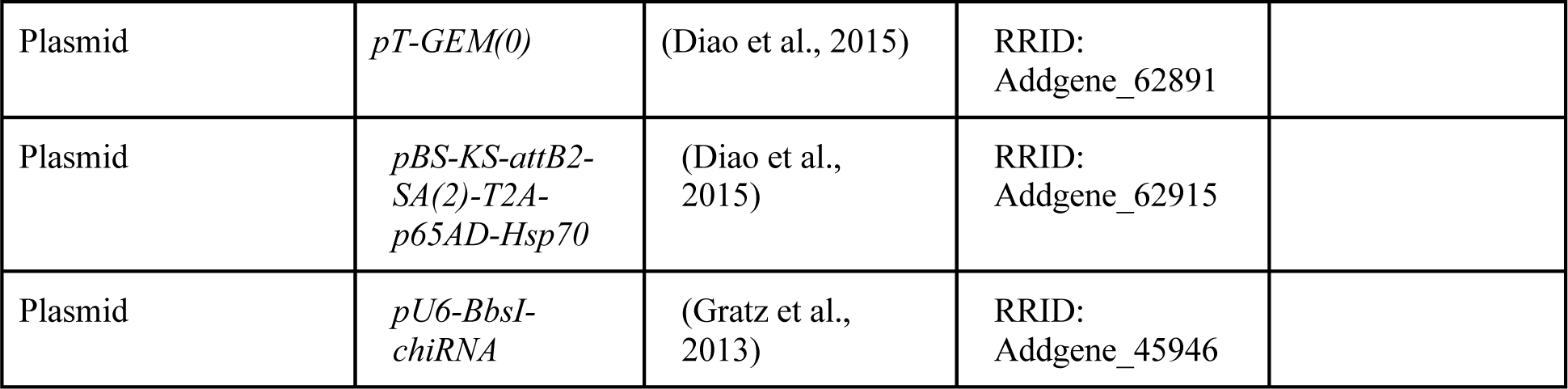

### *Drosophila* Stocks and genotypes

Flies are maintained on standard cornmeal medium at 25 °C with 12-h light–dark cycle. The following lines were used in this study: *GH146-GAL4* (Stocker et al., 1997), *VT033006-GAL4* (Tirian & Dickson, 2017), *Mz19-GAL4* (Jefferis et al., 2004), *knot-GAL4* (Lee et al., 2018), *GH146-Flp, UAS-FRT-STOP-FRT-mCD8-GFP* (Potter et al., 2010), *zfh2-GAL4* (Lee et al., 2018), *Act-FRT-STOP-FRT-GAL4* (Pignoni & Zipursky, 1997), *UAS-Flp* (Duffy et al., 1998), *C15-p65*^*AD*^ (Xie et al., 2019), *DIP-beta-GAL4, DIP-eta-GAL4, DIP-zeta-GAL4* (Carrillo et al., 2015; Cosmanescu et al., 2018), *AstA-GAL4* (Deng et al., 2019), and *elav-GAL4*^*DBD*^ (Luan et al., 2006). *VT033006-GAL4*^*DBD*^, split-GAL4 line #7 (SS01867), #15 (SS01165), and #28 (SS01265) are unpublished reagents generously provided by Yoshi Aso (Janelia Research Campus).

### Generation of *danr-*GAL4^DBD^, kn*-p65*^*AD*^, and *C15-GAL4*^*DBD*^

*danr-GAL4*^*DBD*^ was generated using CRISPR mediated knock-in. ∼2000 bp of genomic sequence flanking the targeted insertion site was amplified by Q5 hot-start high-fidelity DNA polymerase (New England Biolabs) and inserted into *pCR-Blunt-TOPO* vectors (Thermo Fisher). Using this vector, we generated homology directed repair (HDR) vector *TOPO-danr-T2A-GAL4DBD-P3-RFP* by inserting *T2A-GAL4*(*DBD*)::*Zip*+ and *3XP3-RFP-SV40* (cloned from *pT-GEM*(*0*) Addgene #62891) 45bp downstream of the start codon of *danr*. CRISPR guide RNA (gRNA) targeting a sequence inside *danr* (AACATCCGGATGAGCACGCG) were designed by the flyCRISPR Target Finder tool and cloned into a *pU6-BbsI-chiRNA* vector (Addgene #45946). The HDR and gRNA vectors were co-injected into *nos-Cas9* (gift from Dr. Ben White) embryos. RFP+ progenies were selected and individually balanced.

*kn-p65*^*AD*^ was generated by co-injecting *pBS-KS-attB2-SA*(*2*)*-T2A-p65AD-Hsp70* (Addgene #62915) and ΦC31 into the embryos of *MI15480* (BL61064). All *yellow*^*–*^ progenies were individually balanced.

*C15-GAL4*^*DBD*^ was generated using methods similar to *danr-GAL4*^*DBD*^. But because C15 have been shown to be involved in PN dendrite targeting (Li et al., 2017), instead of inserting driver elements into the coding region, the stop codon of *C15* was replaced by *T2A-GAL4*(*DBD*)::*Zip*+ to prevent disruption of the gene.

### Immunofluorescence

Fly brains were dissected and immunostained according to previously described methods (Wu & Luo, 2006). Primary antibodies used in this study included rat anti-Ncad (N-Ex #8; 1:40; Developmental Studies Hybridoma Bank), chicken anti-GFP (1:1000; Aves Labs). Secondary antibodies conjugated to Alexa Fluor 488/647 (Jackson ImmunoResearch) were used at 1:250. 5% normal goat serum in phosphate buffered saline was used for blocking and diluting antibodies. Confocal images were collected with a Zeiss LSM 780 and processed with ImageJ.

### Single-cell RNA sequencing procedure

Single-cell RNA sequencing was performed following previously described protocol (Li et al., 2017). Briefly, *Drosophila* brains with mCD8-GFP labeled cells using specific GAL4 drivers were dissected at appropriate timepoints (0–6h APF, 24–30h APF, 48–54h APF, and 1–5 day adults). Optic lobes were removed from brain during dissection for all timepoints except for 0-6h APF. Single-cell suspension were prepared and GFP positive cells were sorted using Fluorescence Activated Cell Sorting (FACS) into individual wells of 384-well plates containing lysis buffer using SH800 (Sony Biotechnology). Full-length poly(A)-tailed RNA was reverse-transcribed and amplified by PCR following the SMART-seq2 protocol (Picelli et al., 2014). cDNA was digested using lambda exonuclease (New England Biolabs) and then amplified for 25 cycles. Sequencing libraries were prepared from amplified cDNA, pooled, and quantified using BioAnalyser (Agilent). Sequencing was performed using the Novaseq 6000 Sequencing system (Illumina) with 100 paired-end reads and 2 × 8 bp index reads (all except *split#28-GAL4*). *split#28-GAL4* is sequenced using NextSeq 500 Sequencing system (Illumina) with 75 paired-end reads and 2 x 8 bp index reads.

### QUANTIFICATION AND STATISTICAL ANALYSIS

Unless otherwise specified, all data analysis was performed in Python using Scanpy (Wolf et al., 2018), Numpy, Scipy, Pandas, scikit-learn, and custom single-cell RNA-seq modules (Li et al., 2017; Brbić et al., 2020). Gene Ontology analysis were performed using Flymine (Lyne et al., 2007).

### Sequence alignment and preprocessing

Reads were aligned to the *Drosophila melanogaster* genome (r6.10) using STAR (2.5.4) (Dobin et al., 2013). Gene counts were produced using HTseq (0.11.2) with default settings except ‘‘-m intersection-strict’ (Anders et al., 2015). We removed low-quality cells having fewer than 100,000 uniquely mapped reads. To normalize for differences in sequencing depth across individual cells, we rescaled gene counts to counts per million reads (CPM). All analyses were performed after converting gene counts to logarithmic space via the transformation Log2(CPM+1). We further filter out non-neuronal cells by selecting cells with high expression of canonical neuronal genes (*elav, brp, Syt1, nSyb, CadN*, and *mCD8-GFP*). We retained cells expressing at least 8 Log2(CPM+1) for least 2/6 markers.

### Dimensionality reduction and clustering

To select variable genes for dimensionality reduction, we used previously described methods to search for either overdispersed genes (Satija et al., 2015) or ICIM genes (Li et al., 2017). We then further reduced its dimensionality using tSNE to project the reduced gene expression matrix into a two-dimensional space (van der Maaten & Hinton, 2008). We observed that our most recently sequenced cells using NovaSeq (all newly sequenced cells in this study except for *split#28-GAL4*) exhibited some small batch effect with PNs sequenced using NextSeq [*split#28-GAL4*+ PNs and PNs from (Li et al., 2017)]. To overcome this batch effect (in Figure 2, and Figure 7–figure supplement 2 A, C), we performed principal component analysis (PCA) on the ICIM matrix, applied Harmony to correct for batch effect on the principal components (PCs) (Korsunsky et al., 2019), and used tSNE to further project the Harmony-corrected PCs into a two-dimensional space.

To cluster PNs in an unbiased manner, we applied the hierarchical density-based clustering algorithm, HDBSCAN, on the tSNE projection (McInnes et al., 2017). Parameters min_cluster_size and min_samples were adjusted to separate clusters representing different types of PNs. In addition, we also clustered cells using an independent, community-detection method called Leiden on the neighborhood graph computed based on the ICIM gene matrix (Blondel et al., 2008; Levine et al., 2015; McInnes et al., 2018). Both methods appeared to agree with each other for all datasets we examined (examples in Figure 5–figure supplement 1), and we assigned PN types in Figure 2 based on HDBSCAN clustering.

### Global level dynamic gene identification

To identify dynamically expressed genes on the global level (Figure 3), we first identified the top 150 most differentially expressed genes (Mann-Whitney U test) between every two stages and combined them to obtain a set of 474 dynamic genes. We calculated the median expression of each gene at each timepoint and normalized these median expression values by dividing them by the maximum value across time points. We then performed dimensionality reduction on the expression profiles of the genes using tSNE, and identified clusters using HDBSCAN on the projected coordinates. This resulted in identification of 8 sets of genes with distinct dynamic profiles, of which 2 sets are upregulated (Figure 3E), 4 sets are down regulated (Figure 3D), and 2 sets without obvious trend from 0h APF to adult cells (data not shown).

### Transcriptomic similarity calculation

To analyze the transcriptome differences of PNs in different stages (Figure 4E, F), we first isolated lPNs and adPNs to analyze cells from each lineage separately. Cell-level analysis was performed by calculating for each cell mean inverse Euclidean distance in the 2-dimensional UMAP space from all other cells within each stage using the 1215 genes identified by ICIM from most PNs of all stages (Figure 3A). Box plots show the distance distribution at each stage (Figure 4E and F, left). Cluster-level analysis was performed on the MARS clusters. We identified a set of differentially expressed genes for each cluster and calculated Pearson correlation on differentially expressed genes between all pairs of clusters. Bar plots represent mean values across all pairs and errors are 95% confidence intervals determined by bootstrapping with n=1,000 iterations (Figure 4E and F, right).

### PN type identification for most PNs

We observed that the transcriptomes of different PN types are the most distinct at 24h APF and variable genes identified at this stage carry type-specific information (Figure 5). Therefore, we calculated the differentially expressed genes among 24h APF clusters and applied MARS to identify clusters in the space of those genes. MARS is able to reuse annotated single-cell datasets to learn shared low-dimensional space of both annotated and unannotated datasets in which cells are grouped according to their cell types. However, initially we did not have any annotated experiments so we first applied MARS to annotate 24h APF clusters. We then used 24h APF clusters as annotated dataset and moved to annotate PNs at 48h APF. We then repeated the same procedure by gradually increasing our set of annotated datasets. In particular, we used 24h and 48h APF data to help in annotating 0h APF, and finally all three datasets (0h, 24h, 48h) for the adult PNs. We proceed in this order according to the expected difficulty to identify PN types at a particular stage (Figure 5). At each stage, we ran MARS multiple times with different random initializations and architecture parameters to increase our confidence in the discovered clusters, and combined annotations from these different runs. For each cluster, we additionally manually checked the expressions of known PN markers to confirm the annotations.

### Matching clusters representing the same PN type across development using marker expression

For each cluster, we used Mann-Whitney U test to find genes that are highly expressed in that cluster compared to the rest. Then, among those genes, we searched for genes or 2-gene combinations which are uniquely expressed in 1 cluster. We check each gene or combination of genes at the other stages, and if they are also only expressed in 1 cluster and they are of the same lineage, we consider them to be the same types of PNs. Genes used to match clusters representing the same PN types at different timepoints are summarized in a dot-plot in Figure 7–figure supplement 2.

In addition, we used previously sequenced subset of PNs using *Mz19-GAL4* and *kn-GAL4* to overlay with most PNs in combinations of those markers to confirm our matching.

### Matching clusters representing the same PN type across development using similarity calculation

For each cluster, we found the set of differentially expressed genes in that cluster compared to all other clusters at the same stage. Next, we computed the similarity of the sets of identified differentially expressed genes between all pairs of clusters across subsequent stages. Specifically, we computed similarity scores between all pairs of clusters from (i) 0h and 24h APF, (ii) 24h and 48h APF, and (iii) 48h and adult APF. The similarity of the sets of differentially expressed genes was computed as the Jaccard similarity index defined as the ratio of the cardinality of the intersection of two sets and the cardinality of the union of the sets. We excluded clusters representing vPNs and APLs for matching most PNs across 4 stages (Figure 7). For each cluster, we then identified its most similar cluster at the adjacent stage according to the Jaccard index. If the clusters between two stages coincide—meaning that two clusters from two stages have the highest similarity to each other, we consider the clusters to be matched. Empirically, we found this matching procedure to be stringent, resulting in high confidence matching pairs.

### Identification of type specific dynamic genes

We first identified all dynamic and stable genes. For each PN type matched across all developmental stages, we consider all genes that significantly change their expression between any two adjacent time points as dynamic genes. Statistical significance was determined by two-tailed t-test and p-values were adjusted for multiple hypothesis testing using Benjamini–Hochberg method. To ensure that gene is expressed in at least one time point, we required that Log2(CPM+1) > 2 in at least 50% of cells at one time point. Further, for each PN type we characterized all genes with FDR adjusted p-value larger than 0.9 at all time points as stable genes.

If the same gene is identified as dynamic in two PN types at the same time transition but it shows opposite dynamics, we consider it as dynamic-dynamic gene (Figure 8A). Here, opposite dynamics means that the mean expression increases in one PN type in the transition from one stage to another but decreases in the another PN type. On the other hand, if the same gene is identified as dynamic in one PN type but stable in another PN type, we consider it as dynamic-stable gene (Figure 8A).

### Correlation between different PN types

MARS clusters of excitatory PNs were used for analysis in Figure 9. We performed PCA on the entire matrix and calculated their correlation based on the PCs. Dendrograms shown in Figure 9– figure supplement 1 are generated using distance calculated using Farthest Point Algorithm and organized so the distance between successive leaves is minimal.

To observe the relationship between birth timing and their transcriptomic similarity, for each stage, we selected adPN clusters, performed PCA among all genes detected, calculated their correlation, and plotted the correlation matrices according to their birth order (Yu et al., 2010) (Figure 9B). For the two clusters representing either VM7 or VM5v PNs, we ordered them based on their correlation with decoded PN types whose birth order are adjacent to either of these two PN types. We are showing adPNs in the figure because we decoded much fewer transcriptomic clusters belonging to the lPN lineage, which is too few to carry out analysis shown in Figure 9 C– D with robust statistical backing. Nevertheless, we still observed higher correlation between lPN types with adjacent birth-order in 0h and 24h APF (data not shown).

### Spearman’s rank correlation calculation and permutation test

For consistency, 8 adPN types that were decoded across 4 stages were selected for this analysis (Figure 9C). For each PN type X, the group of PNs that are born either earlier or later than X was selected depending on which direction contains more PN types (each group contains at least 5 types of PNs). Then, we ranked the PN types according to their correlation with X and calculated the Spearman’s rank correlation of this ranking with the ranking based on their birth order. For each stage, we obtained the average correlation coefficients and plotted the result as a red dot on the x-axis for each timepoint. Higher value indicates higher correlation between birth order and order calculated based on their transcriptomic similarity.

To determine if we can reject the null hypothesis that the adPN transcriptomic similarity do not covary with the ranks of the birth order, we performed permutation test. We randomly shuffled the birth order and performed the aforementioned correlation calculation for 5000 iterations. The distribution of the simulated average correlations is shown in the histogram of Figure 9C. We obtained the p-value by dividing the number of times of the simulated correlation is greater than the observed correlation by the total number of iterations.

### Developmental trajectory analysis

Pseudo-time analysis of 0h APF adPNs was performed using the monocle package in R (Trapnell et al., 2014; Qiu et al., 2017; Cao et al., 2019). We selected only adPNs born at larval stage because the embryonically born adPNs have a very distinct transcriptomes which skew clustering. We applied the dimensionality reduction method UMAP (Becht et al., 2018) on 561 24h ICIM genes to resolve distinct PN types. This dimensionally reduced dataset was then used as the basis for a developmental trajectory graph created by Monocle 3. We then selected the cluster representing DL1 PNs to be the root node of the trajectory and computed the pseudo-times based on distance from the root in accordance to the trajectory.

## Acknowledgement

We thank Yoshi Aso, Gerald Rubin, Hugo Bellen, Kai Zinn, and Larry Zipursky for the kind gifts of reagents. We thank the Bloomington *Drosophila* Stock Center and the Vienna *Drosophila* Resource Center for fly lines, and Addgene for plasmids. We thank Tom Clandinin, Yanyang Ge, Julia Kaltschmidt, Justus Kebschull, Kang Shen, Andrew Shuster, and all Luo lab members for technical support and insightful advice on this study. We thank Mary Molacavage for administrative assistance.

## Additional information

### Author contributions

Qijing Xie, Conceptualization, Methodology, Software, Validation, Formal Analysis, Investigation, Resources, Data Curation, Writing–Original Draft, Writing–Review & Editing, Visualization; Maria Brbic, Methodology, Software, Formal Analysis, Resources, Data Curation, Writing–Review & Editing, Visualization; Felix Horns, Resources; Sai Saroja Kolluru, Resources; Bob Jones, Resources; Jiefu Li, Resources; Anay Reddy, Resources; Anthony Xie, Formal Analysis; Sayeh Kohani, Formal Analysis; Zhuoran Li, Resources; Colleen McLaughlin, Resources; Tongchao Li, Resources; Chuanyun Xu, Resources; David Vacek, Resources; David J. Luginbuhl, Resources; Jure Leskovec, Resources; Stephen R. Quake, Resources, Funding Acquisition; Liqun Luo, Conceptualization, Resources, Writing–Original Draft, Writing–Review & Editing, Supervision, Funding Acquisition; Hongjie Li, Conceptualization, Methodology, Formal Analysis, Investigation, Resources, Data Curation, Writing–Review & Editing, Supervision.

---

**Figure 1—figure supplement 1.**
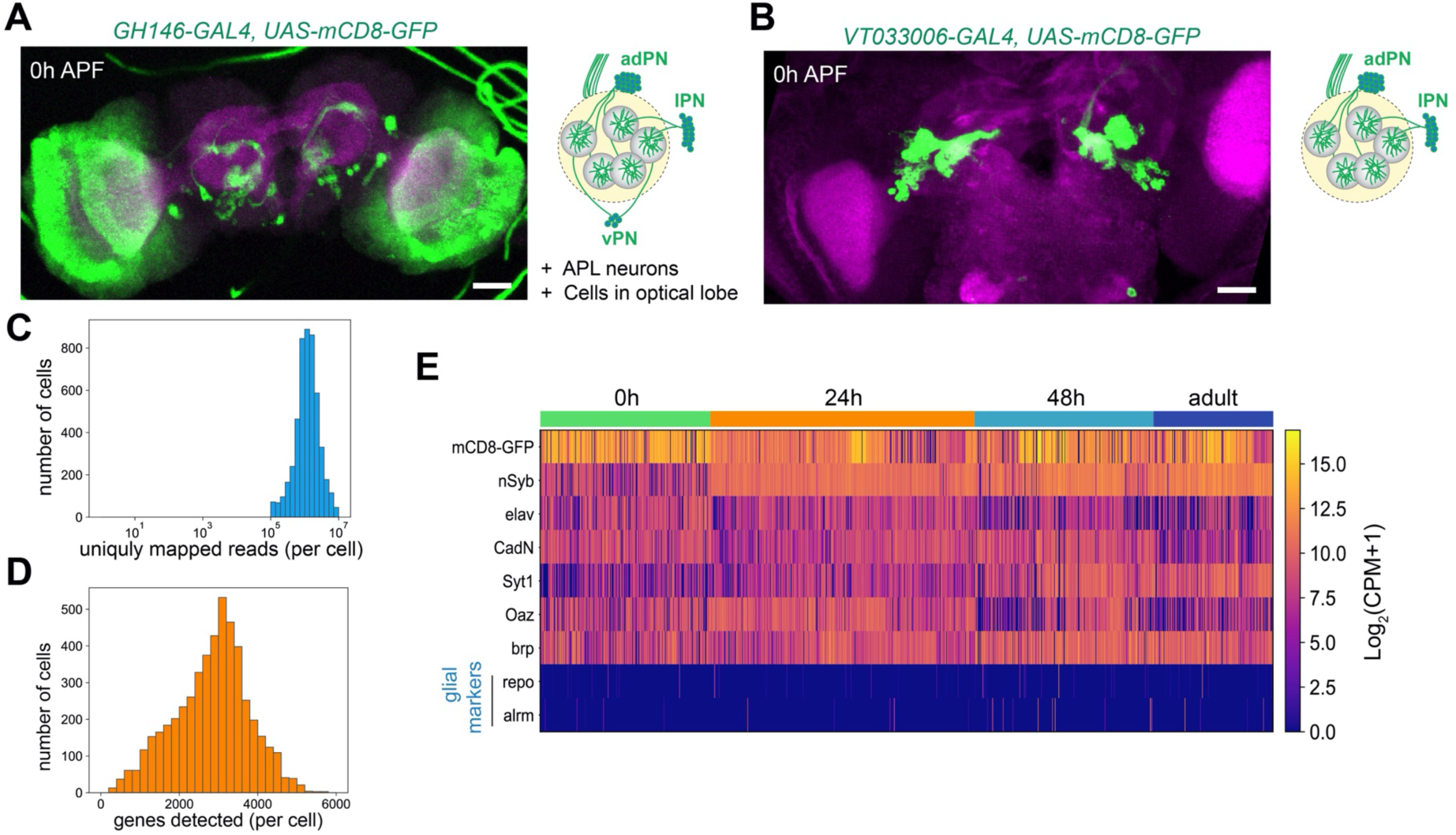
Technical characteristics of PN scRNA-seq. **(A)** Representative confocal image and illustration of cells labeled by *GH146-GAL4* at 0h APF. Other than PNs and a pair of APL neurons in the central brain, many cells in the optic lobes are also labeled. **(B)** Representative confocal image and illustration of cells labeled by *VT033006-GAL4* at 0h APF. This driver labels excitatory PNs, but not cells in the optic lobes or vPN or APL neurons. Scale bars, 40 μm. **(C)** Distribution of the number of uniquely mapped reads per cell. **(D)** Distribution of the number of detected genes per cell. **(E)** Heatmaps showing the expression of: *mCD8-GFP*, pan-neuronal makers (*nSyb, elav, CadN, Syt1*, and *brp*), PN marker (*Oaz*), and glial markers (*repo* and *alrm*). Expression levels are indicated by the color bar (CPM, counts per million).

**Figure 2—figure supplement 1.**
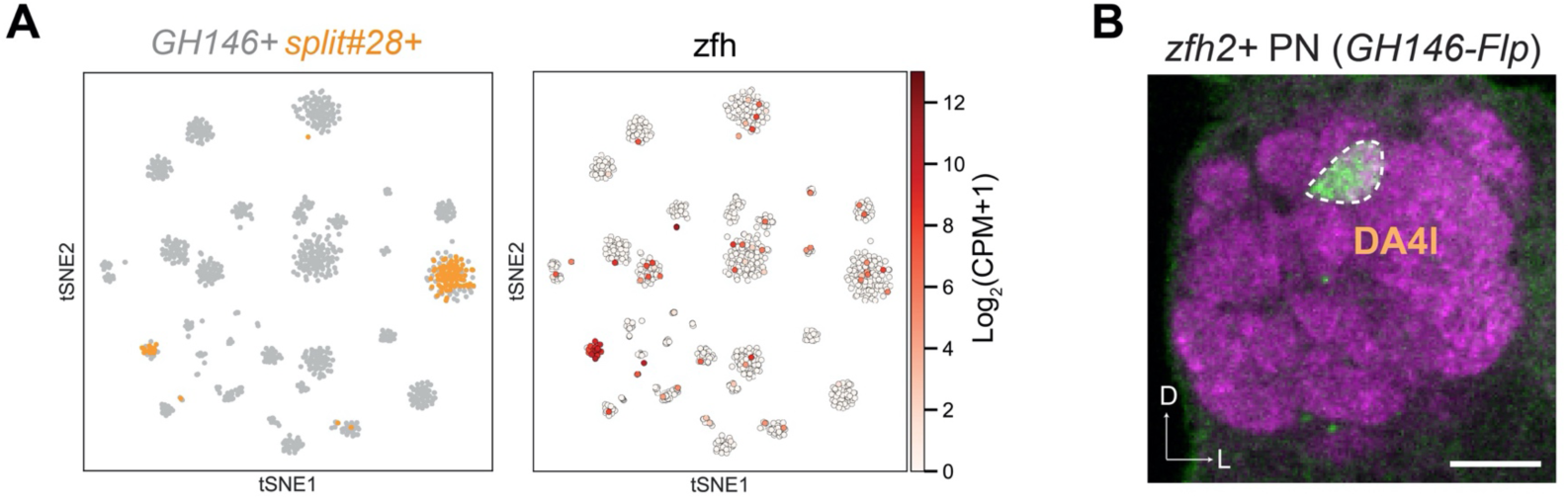
Validation of DA4l PN identity. **(A)** Visualization of *GH146*+ and *split#28-GAL4*+ PNs using tSNE. Cells are colored according to driver genotypes (left) or by the expression of *zfh2* (right). **(B)** *zfh2-GAL4*, after intersecting with *GH146-Flp*, labels DA4l PNs. Scale bars, 20 μm. Axes, D (dorsal), L (lateral).

**Figure 2—figure supplement 2.**
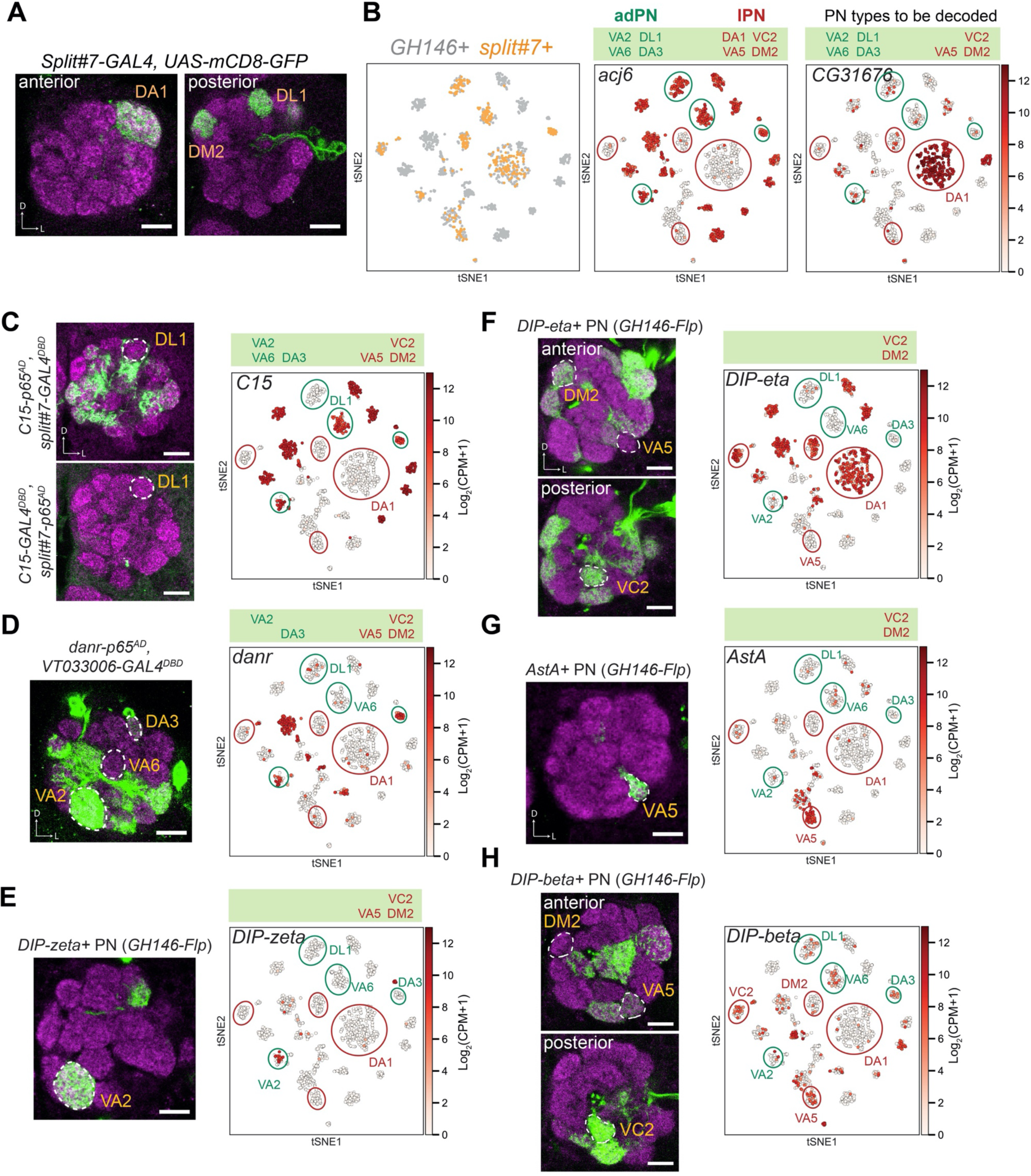
Decoding *split#7*+ PNs. **(A)** Representative confocal images of *split#7*+ PNs. Without permanent labeling, this driver is strongly expressed in 3 PN types in adults. Permanent labeling showed that it can label 8 adult PN types (Figure 2D), suggesting that this driver is expressed in 8 PN types during development and turned off in 5 of them in adult stage. **(B)** Visualization of *GH146*+ and *split#7*+ PNs colored according to genotype (left), *acj6* (middle), and *CG31676* (right) expression. Previously, we know among those *split#7*+ PNs, the cells with *CG31676* expression are DA1 PNs (Li et al. 2017). **(C)** Among *split#7*+ adPN clusters (circled in green), only one cluster does not express *C15*. Intersection between *C15-p65*^*AD*^ and the GAL4 DNA-binding domain (DBD) from *split#7* (top) as well as intersection between *C15-GAL4*^*DBD*^ and the p65-activating domain (AD) from *split#7* (bottom) revealed that the *C15* negative cluster represents DL1 PNs. **(D)** Among *split#7*+ adPNs (circled in green), two clusters are *danr–*. One of those cluster represents DL1 PNs. Intersection between *danr-GAL4*^*AD*^ and *VT033006-GAL4*^*DBD*^ (split-GAL4 with PN specific expression) revealed the other *danr–* adPN is VA6 PNs. **(E)** One *split#7*+ cluster specifically expresses *DIP-zeta*. Intersection between *DIP-zeta-GAL4* and *GH146-Flp* revealed this cluster represents VA2 PNs. As three out of four adPN clusters are assigned, we assigned the last unassigned to be DA3 PNs. **(F)** Among *split#7*+ lPNs (circled in red), only one cluster is *DIP-eta*+. Intersection between *DIP-eta-GAL4* and *GH146-Flp* revealed the identity of this cluster as VA5 PNs. **(G)** The *DIP-eta-*cluster also specifically expresses *AstA*. Intersection between *AstA-GAL4* and *GH146-Flp* only labels VA5 PNs, further confirming its identity. **(H)** Among the last two unmapped clusters, one is *DIP-beta*+. Intersection between *DIP-beta-GAL4* and *GH146-Flp* revealed the cluster negative for *DIP-beta* is DM2 PNs. And we assigned the remaining *split#7*+ lPN cluster to be VC2 PNs. Scale bars, 20 μm. Axes, D (dorsal), L (lateral).

**Figure 2—figure supplement 3.**
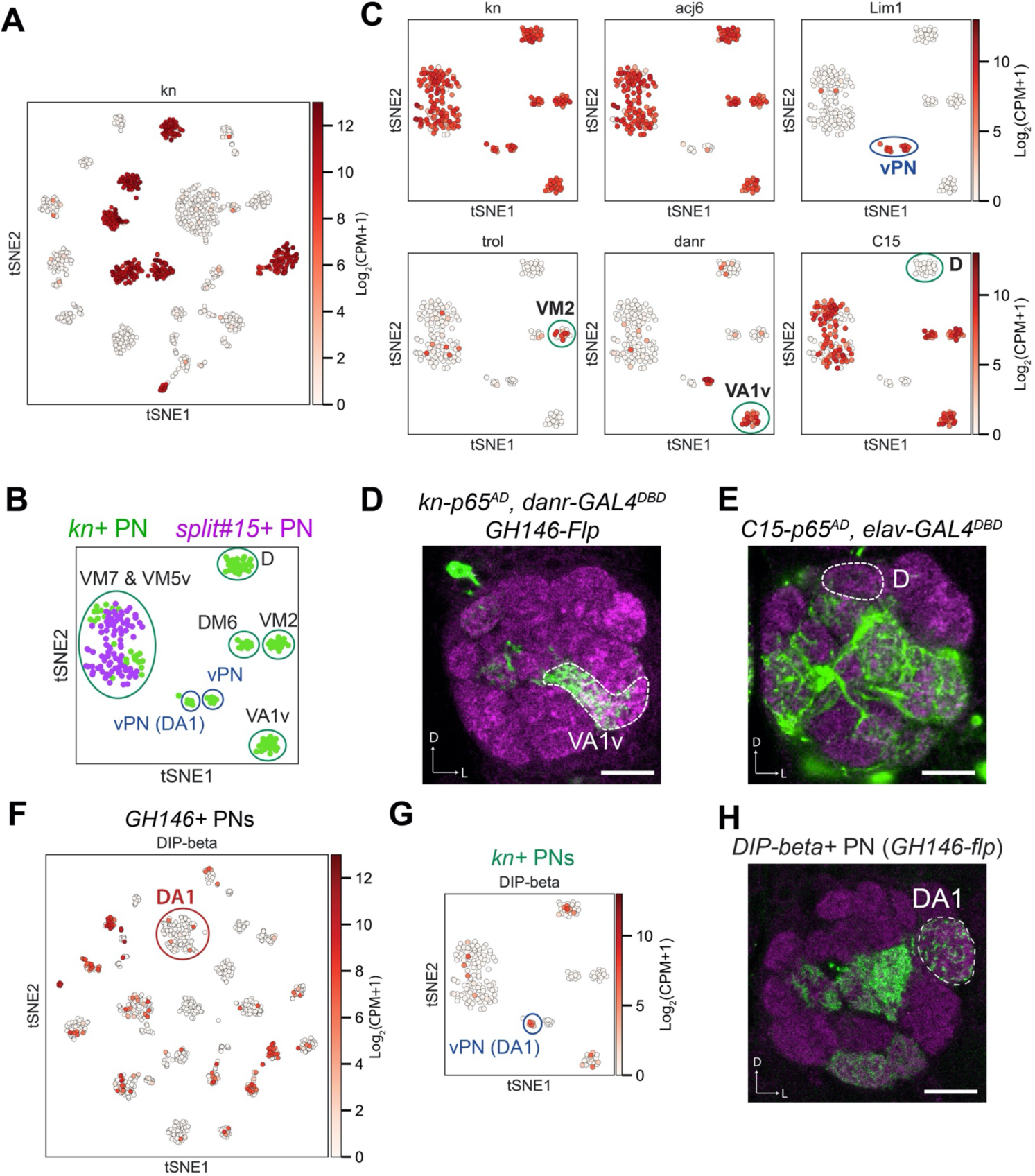
Decoding the identity of *kn*+ PNs. **(A)** *kn* is expressed in 7 transcriptomic cluster in *GH146*+ PNs at 24h APF. **(B)** Visualization of *kn*+ and *split#15-GAL4*+ PNs at 24h APF using tSNE. *kn*+ PNs (green) form 8 clusters, two of them intermingled with *split#15*-*GAL4*+ PNs (purple). These 8 clusters are assigned to specific PN types using information in the following panels. **(C)** Summary of marker genes used to decode the identity of *kn-GAL4*+ PNs. *trol*+ cluster represents VM2 PNs (Li et al., 2007). **(D)** Intersection between *kn-p65*^*AD*^ and *danr-GAL4*^*DBD*^ with *GH146-Flp* revealed that the cluster positive for both *kn* and *danr* is VA1v PNs. **(E)** Intersection between *C15-p65*^*AD*^ and *elav-GAL4*^*DBD*^ revealed that the cluster positive for *acj6* but negative for *C15* is D PNs. **(F)** Visualization of *DIP-beta* expression among *GH146*+ PNs. DA1 lPNs does not express *DIP-beta*. **(G)** Visualization of *DIP-beta* expression among *kn*+ PNs. One vPN cluster expresses *DIP-beta*. **(H)** Representative confocal image of *DIP-beta-GAL4* after intersecting with *GH146-Flp*. Innervation of the DA1 glomerulus indicated the *DIP-beta*+ vPN cluster is vPN (DA1). Scale bars, 20 μm. Axes, D (dorsal), L (lateral).

**Figure 4—figure supplement 1.**
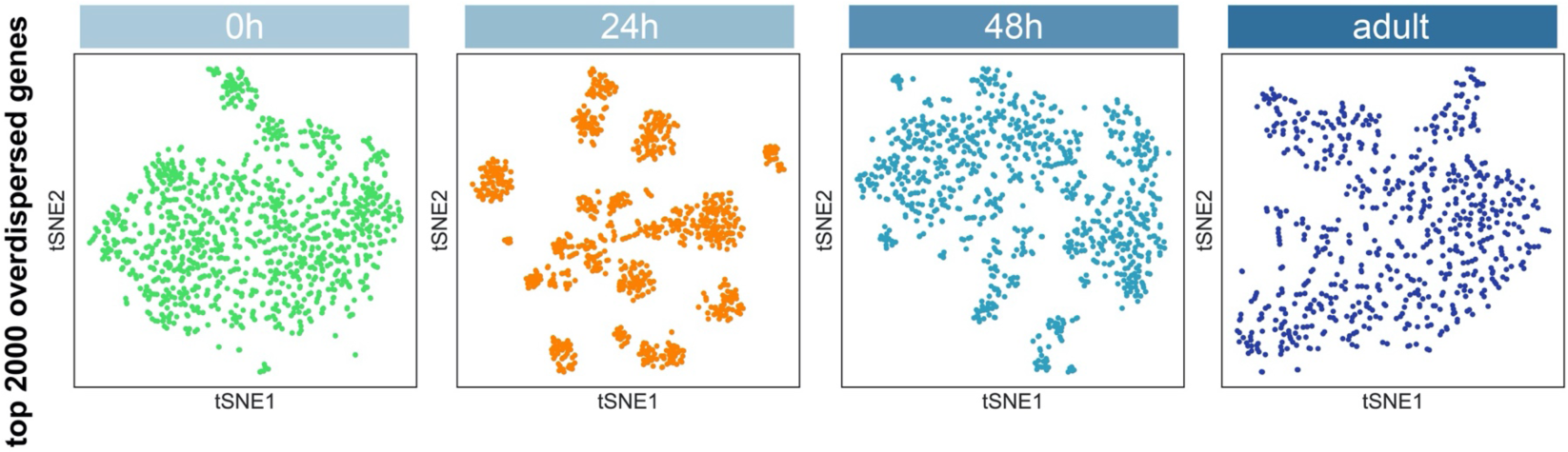
Visualization of most PNs at different stages using tSNE. Dimensionality reduction was computed using overdispersed genes found at each stage.

**Figure 4—figure supplement 2.**
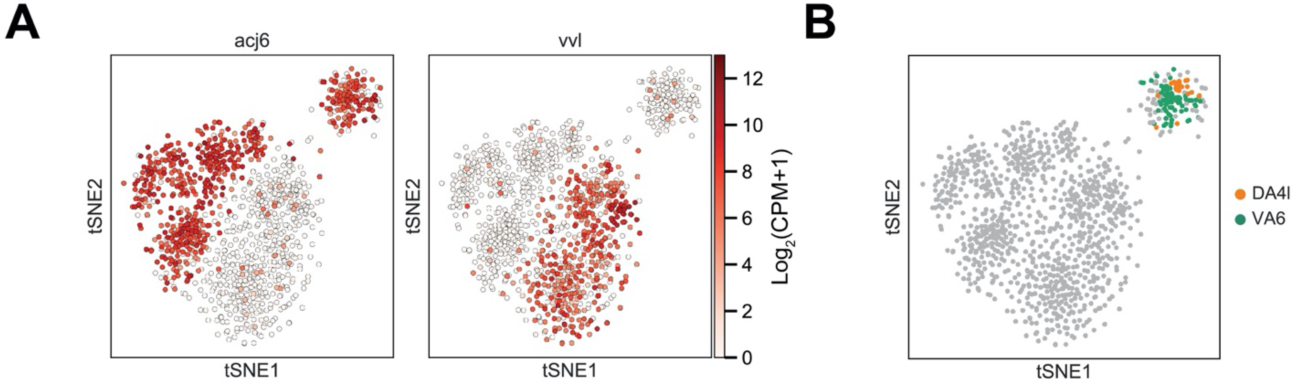
Embryonically born and larval born PNs at 0h APF. **(A)** The larger cluster consists of both adPNs (*acj6*+) and lPNs (*vvl*+) while the smaller cluster contains only adPNs. **(B)** Two types of embryonically born PNs, DA4l and VA6 PNs, are both mapped to the smaller cluster (details in Figure 7).

**Figure 5—figure supplement 1.**
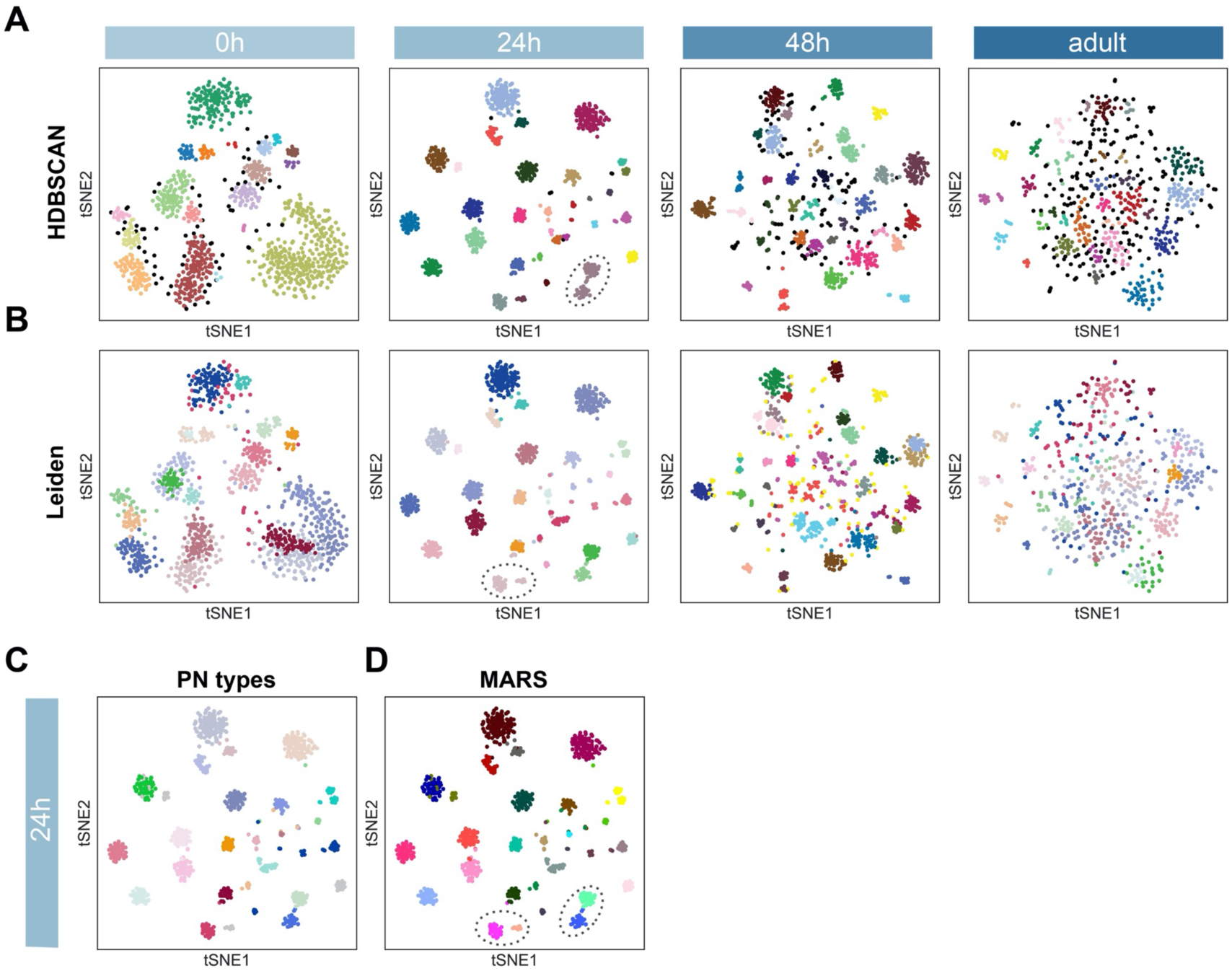
PN type identification using two other independent methods. **(A)** Dimensionality reduction by 24h ICIM genes followed by cluster identification using HDBSCAN. Circled cells belong to two PN types but are assigned to the same cluster using HDBSCAN. **(B)** Cluster identification by Leiden based on neighborhood graph computed on 24h ICIM genes. Circled cells belong to two PN types but are assigned to the same cluster using Leiden. **(C)** 24h APF PNs colored according to PN types validated in Figure 2. **(D)** PN types identified using MARS (same as Figure 5B). PN types which are incorrectly annotated by HDBSCAN or Leiden are correctly annotated as distinct clusters by MARS.

**Figure 6—figure supplement 1.**
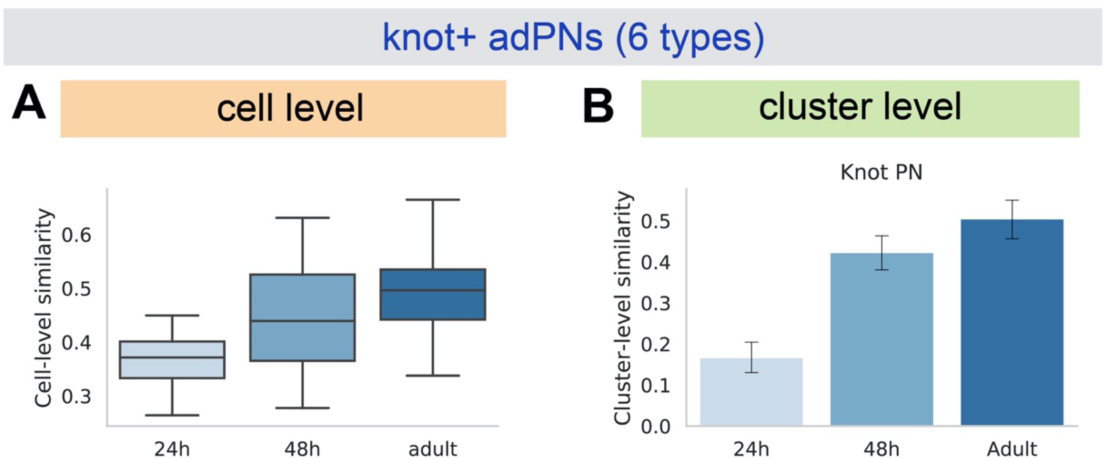
*kn*+ adPN transcriptomes become more similar as development proceeds. **(A)** Box plot of Euclidean distance between all pairs of *kn*+ cells using ICIM genes identified among them. *kn*+ vPNs are excluded from this analysis. 24h APF: 98 cells, mean ± standard deviation: 0.374 ± 0.066; 48h APF: 174 cells, mean ± standard deviation (std): 0.446 ± 0.912; adult: 124 cells, mean ± std: 0.493 ± 0.085 **(B)** Bar plot of Pearson’s correlation between all pairs of *kn*+ adPN clusters. 24h APF: 8 clusters, mean ± std: 0.167 ± 0.141; 48h APF: 8 clusters, mean ± std: 0.424 ± 0.170; adult: 8 clusters, mean ± std: 0.506 ± 0.187.

**Figure 7—figure supplement 1.**
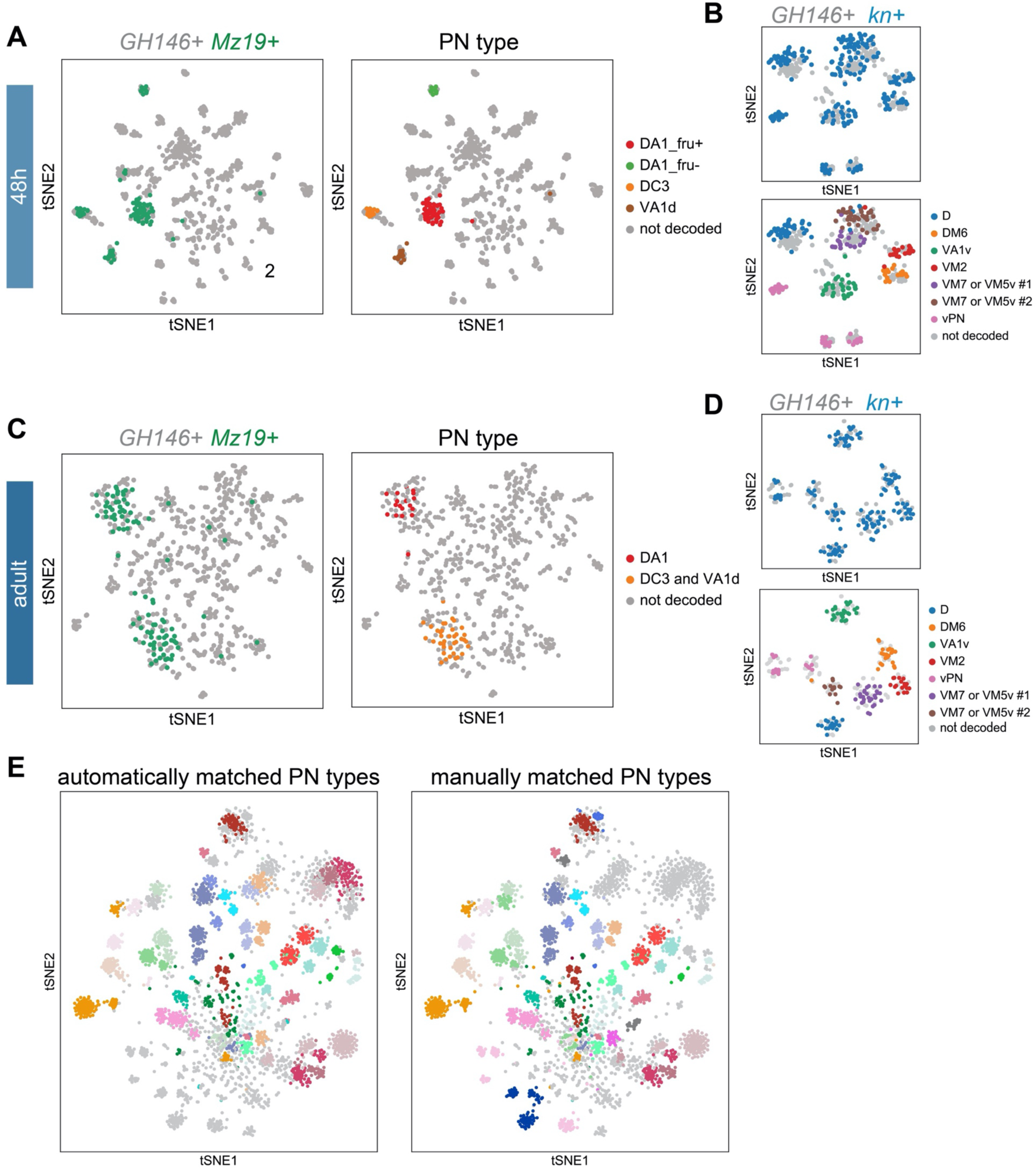
Supporting evidence for matching PN types across developmental stages. **(A, C)** Visualization of sequenced *GH146*+ PNs (grey) with *Mz19*+ PNs (green) at 48h APF (A) and at the adult stage (C). PN type of *Mz19*+ PNs shown on left were decoded previously (Li et al. 2017). **(B, D)** Visualization of *kn*+ PNs from cells sequenced using *GH146-GAL4* (in grey) and cells sequenced using *kn-GAL4* (in blue) at 48h APF (A) and at the adult stage (C). Annotation of *kn-GAL4*+ cells was done in Figure 6. **(E)** Visualization of the same types of PNs matched automatically or manually. Transcriptomic clusters representing the same PN types of different developmental stages are labeled in the same color. Colors used to indicate PN types are consistent with those in Figure 7B.

**Figure 7—figure supplement 2.**
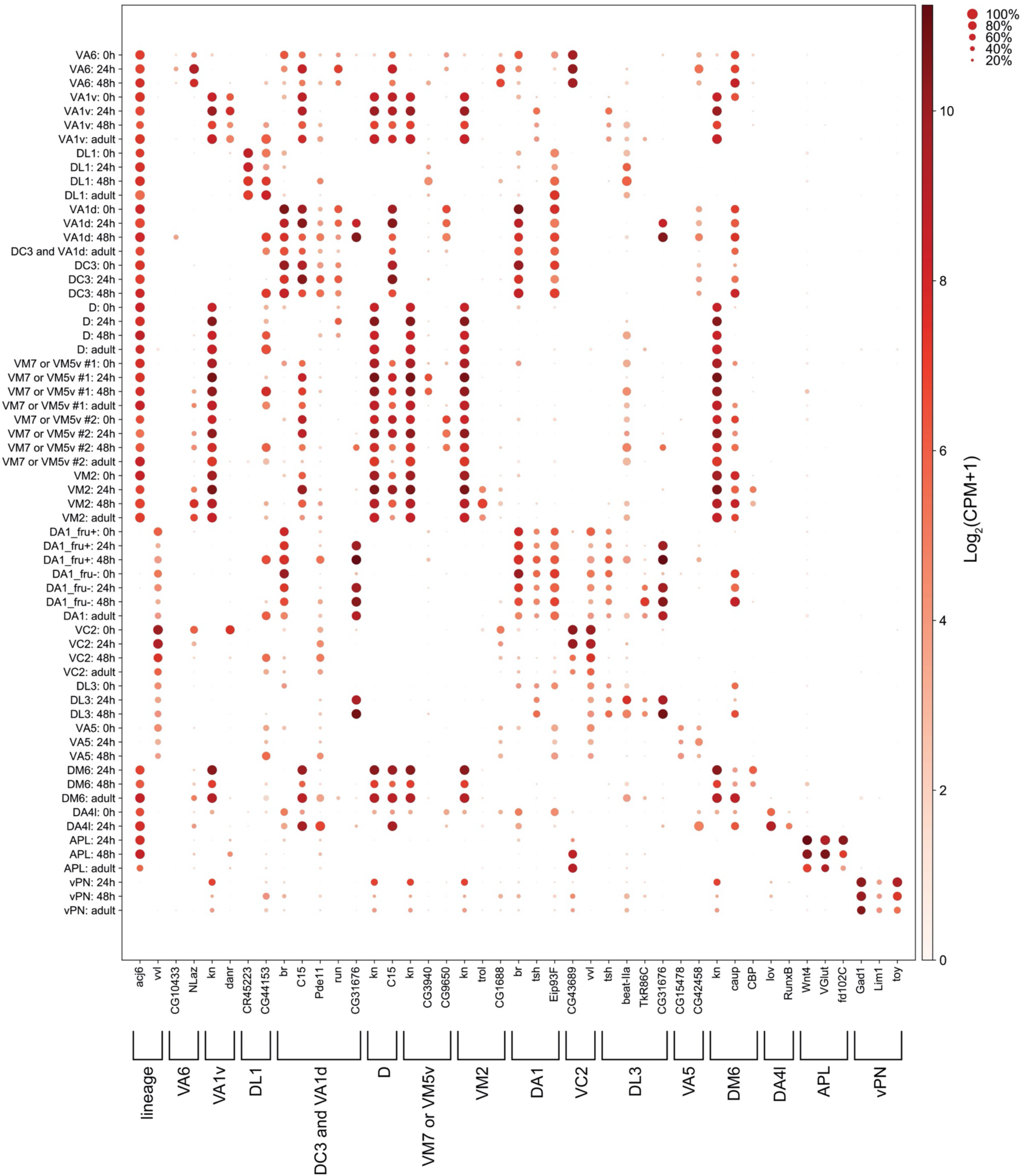
Markers used for manually matching PNs. Dot plot of markers used to match the same types of PNs across different stages. Size of the dot represents percentage of cells expressing a given marker in a cluster at a given stage, and color of the dot represents expression level.

**Figure 9—figure supplement 1.**
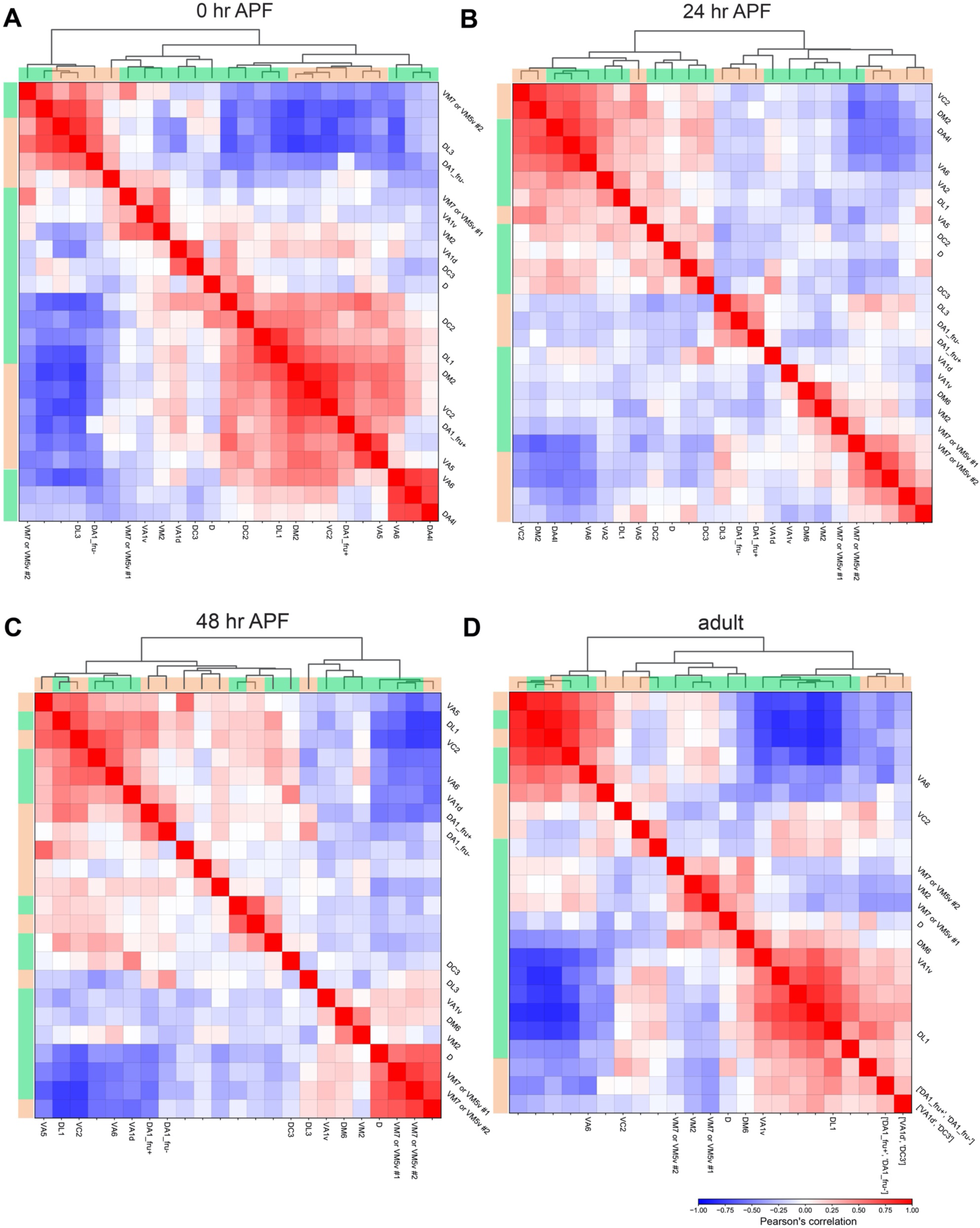
Hierarchical clustering of all excitatory PNs. Hierarchical clustering of all excitatory PNs of 0h APF **(A)**, 24h APF **(B)**, 48h APF **(C)**, and adult **(D)**. Correlation calculation and hierarchical clustering is done on the principal components calculated using the entire gene matrix. adPNs are indicated by green bar and lPNs are indicated by orange bar on the top and left side of each plot. Clusters that have been matched to specific PN types are labeled.

